# Retrieval uncertainty drives rapid consolidation of confabulated memories in retrosplenial cortex

**DOI:** 10.64898/2026.09.01.748171

**Authors:** Aditya Singh, Nirupam Das, Palash Ahuja, Aayush Banerjee, Srujana Kaluve, Aparajita Asane, Sankhanava Kundu, Titli Saha, Sugandhita Rajesh, Balaji Jayaprakash

## Abstract

Learning to make associations in the absence of direct reinforcement, based on previous memories of reward or punishment, is an evolutionary trait quintessential for survival. Little is known about if such higher order associations (HOA) can lead to fabrication of new memories for unreal events. We probed if uncertainty during retrieval of old memory representations can trigger such confabulation. Using chemogenetics, *in vivo* imaging, and behavior we show that uncertainty in memory perception can not only transfer conditioning to hitherto unexperienced novel context but also can get rapidly consolidated. We find while acquisition of HOA require dorsal-CA1 parvalbumin interneuron activity in intact hippocampus, retrieval requires retrosplenial cortex (RSc). We develop an automated **La**rge-scale **B**rainwide **C**orrelated **A**ctivity **M**apping (LaBCAM) method to identify functional connectivity changes across different brain regions in such scenarios. LaBCAM finds that only uncertain remote retrieval engages canonical fear circuitry while regular retrieval does not. Longitudinal imaging of dendritic spines in RSc, a region involved in detection and context-dependent conflict resolution, reveals differential reorganization of spines during acquisition and rapid systems consolidation of HOA. Interestingly, we find HOA is implicated early in AD mice model (APP/PS1) and also in aged animals.

## Introduction

Perils of basing judgements solely on eye-witness testimony have been investigated extensively^1–3^. Malleable nature of memories has led to several instances of prosecuting innocent suspects^4–6^. Little is known about the effect and underlying neural mechanisms of lining up innocents along with suspects for identification parades, on the victim’s memory of the incident and the perpetrator^7,8^. Given the distributed nature of long-term systems consolidated memory^9–13^, we hypothesize that, the memory itself is deconstructed into its feature elements and any higher order association formed using these elements could result in a false memory percept of hitherto unexperienced life event. Here we test if this reconstructive nature of remote memory recall can lead to development of confabulated memory for a previously unseen, novel context.

Acquisition of novel information from everyday experiences requires the hippocampus^14^. During a learning episode, the identity of the features experienced through different sensory modalities is thought to be integrated via hippocampus as context representations^15–20^. Such integration is specific and comprises of various features and elements^21,22^. This initial encoding undergoes systems consolidation (SysCon) when statistical regularities across various features or elements of configural representations are extracted and stored for the long-term elsewhere in the cortex ^23–29^. During SysCon, some information about the contextual elements is lost, leading to generalized, less-detailed (coarse) remote retrieval^30–32^. However, studies on long-term cued recall suggest that even those missing details of remote memories can be recalled when cued^33–38^. Also, we have previously shown that the content of some feature elements is remembered for a long time while other features are forgotten^39^, consistent with the prevailing idea that the representation of generalized memories is elemental. Here we probe if such deconstructed memories can serve as a substrate for higher order associations.

Real-world learning involves forming associations, when a response is encoded for novel stimuli based on previous experiences (rewarding or alarming, unconditioned stimulus; US). Whereas, experiential learning often requires development of association between two neutral stimuli^40–44^ - CS1(preconditioned) and CS2(new) in the absence of US through secondary reinforcement^45–48^. Such higher order association (HOA) is an evolutionarily conserved vital learning mechanism observed across various species^49–52^. All the previous studies on higher order conditioning have focused on associations developing within a short delay of first-order learning, ranging from several minutes to hours^53–57^. There is a gap in knowledge regarding the neural and behavioral mechanisms underlying the acquisition of HOA during remote memory retrieval in healthy, aging, and diseased conditions such as Alzheimer’s disease (AD).

## Results

### Uncertainty during remote retrieval fabricates memory through HOA

Memory for training context, when retrieved remotely, tends to generalize across related contexts^30,31,58–61^. These generalized representations lack the specific details when retrieved remotely. This apparent lack of precision makes the definition of the training context **A** fuzzy, resulting in uncertainty during remote retrieval of this memory in a similar context **B**^11,25,39,58^. We designed experiments to investigate if these uncertainties can lead to creation of false belief through HOA for a perceived event that never occurred^44,62^. Specifically, using contextual fear conditioning we probed if mere retrieval of an old memory (training in context **A**) in an uncertain context **B** that has some similarity to the training context, can lead to development of fear for a novel context **C** that is *distinct* from training context **A**, and if retrieving order plays a role in such false memories.

Figure 1a (Venn diagram) summarizes the relationship between contexts used where we establish the relationships among the contexts through a combination of discriminatory and recent memory testing (see methods, Supplementary Material, Figure M1, M2). Our premise is that during remote testing of the retrieval of fear memory in the training context **A**, in itself being a recent experience, would result in the initial hippocampal encoding. When this salient event is followed by an uncertain event the next day, such as retrieval in a similar context **B**, the unique defining features of context **B** that were neutral until now can acquire the fear associated with recent fearful event through its common elements that overlaps with definition of original training context **A**. Such fear transfer can result in development of fear for a very distinct context (context **C**) even though it shares its features only with **B** (uncertain context) and not with **A** (original training context).

**Figure 1.**
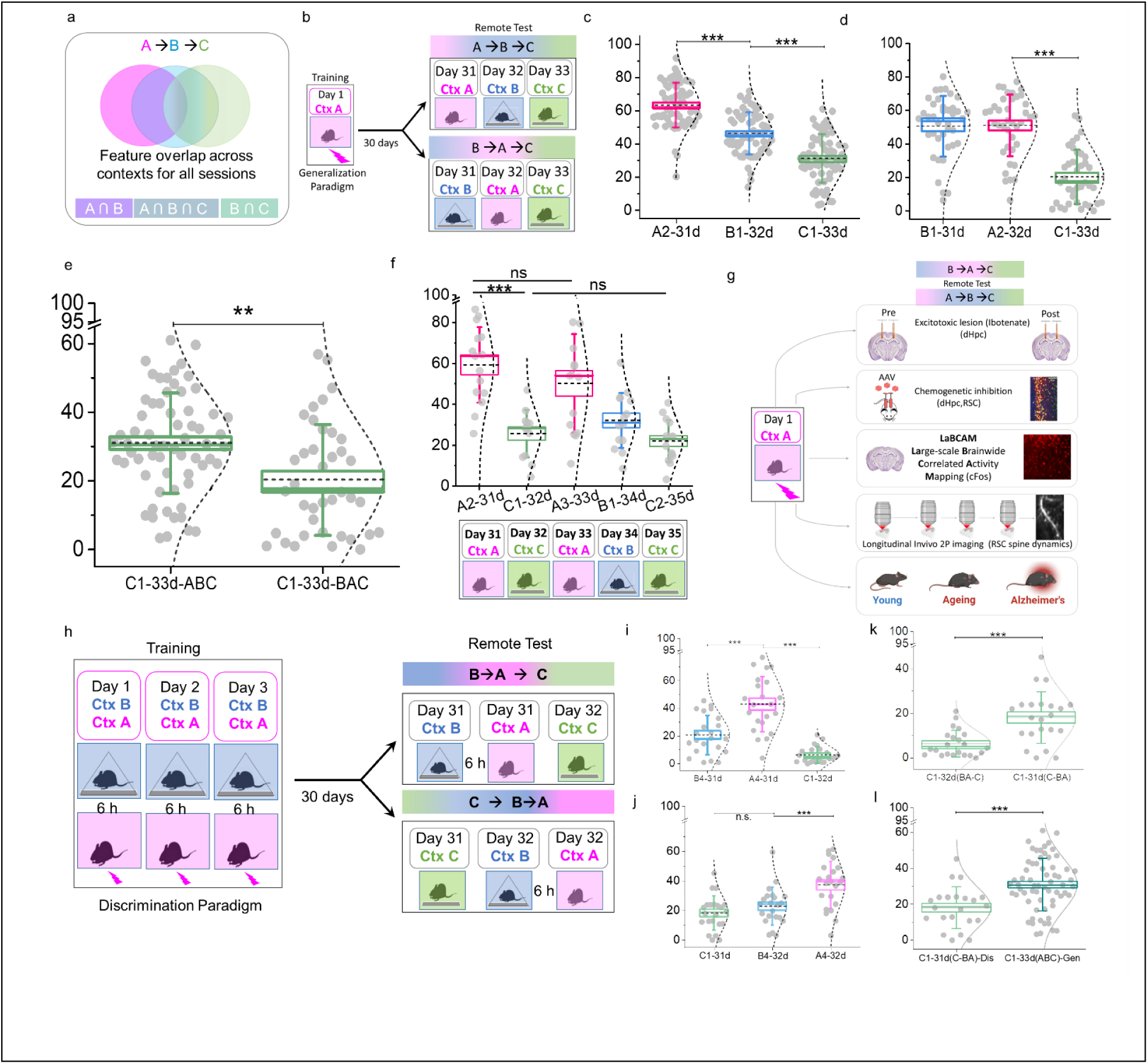
Fear fabrication during remote retrieval. Animals show characteristic memory retrieval across contexts A, B, and C for both generalization and discrimination paradigms. a) Venn diagram shows the feature overlap across the three contexts. b) Animals are trained in context A and same animals are sequentially tested either in ABC or BAC order after a month of one-shot fear learning. During training in generalization paradigm, 120sec initial exposure is followed by three 2sec long shocks at 30sec interval each. Animals are exposed to respective context for 120-150sec during all tests. c, d, e) Testing order dependent remote memory retrieval and higher order associations through second order conditioning (SOC). Animals (n = 117) are trained and tested in generalization paradigm (b). c) ABC-subgroup tested at 24-hour interval in A, B, and C on days 31, 32, and 33 after training, shows context discrimination with decreasing fear (Freezing in A(mean±sem): 63.22±1.68, B: 46.08±1.58, C: 30.99±1.83; one-way, repeated measures or 1-w RM ANOVA, F_2,62_ = 153.34, P>F = 9.95E-25, n = 64). Consistent with the trend of decreasing feature overlap across the three contexts, freezing in A was significantly higher than B (|t|_126_ = 10.32, P > |t| = 5.45E-18) and C (|t|_126_ = 19.40, P > |t| = 3.59E-39). Freezing in B was also significantly higher than C (|t|_126_ = 9.08, P > |t| = 5.68E-15). d) BAC-subgroup tested at 24hour intervals in B, A, and C on days 31, 32, and 33 after training showed overall significantly different freezing across the contexts (Freezing in B: 50.52±2.90, A: 51.02±2.94, C: 20.24±2.59; 1-w RM ANOVA, F_2,37_ = 62.18, P>F = 1.47E-12, n = 39). In contrast to ABC-group, we observe testing order dependent context generalization between similar context B and training context A (|t|_76_ = 0.25, P > |t| not significant). The effect of testing order supersedes feature overlap for B and A but not for C as animals can discriminate the different context C from both B (|t|_76_ = 10.75, P > |t|= 1.94E-16) and A (|t|_76_ = 10.93, P > |t|= 9.03E-17). During ABC-testing order, we hypothesize second order conditioning based higher order associations develop between the overlapping and non-overlapping feature representations of context B and A. Such remote retrieval dependent new learning leads to consistently higher fear recall for context C specifically in ABC testing order(e), as C has higher feature overlap with B but minimal feature overlap with A. e) Freezing in context C was significantly higher in the ABC cohort than in the BAC cohort (n = 64, AB**<u>C</u>**: 30.99 ± 1.83 vs. n = 39, BA**<u>C</u>**: 20.24 ± 2.59; F_(1, 101)_ = 12.09, P>F = 7.50E-4), indicating that prior retrieval in the training context A renders subsequent exposure to the related context B uncertain, a pre-requisite for HOA. f) After training in generalization paradigm, when remote fear memory is tested in A on day 31 and C on 32 without intervening exposure to B, the minimal feature overlap is insufficient for developing fear-based higher order associations as indicated by low freezing in context C-32d. Animals show different freezing across contexts (1w-RM ANOVA, F_(4, 9)_ =18.89, P>F=2.1E-4). Freezing in A2-31d (56.56 ±5.05) and A3-33d (44.59±5.64) is relatively higher than C1-32d (25.62 ±3.22), B1-34d (28.89 ±3), and C2-35d (20.49±2.62). Statistically, freezing was similar for A2-31d and A3-33d (Post hoc Tukey’s test: ltl_48_=3.46, P>|t|=not significant), while animals could distinguish context A-31d from C1-32d (|t|_48_=8.96, P>|t|=8.0E-7). Interestingly, animals showed similar freezing in C1-32d and C2-35d (post hoc Tukey’s test: |t|_48_=1.48, P>ltl=not significant), indicating that prior exposure to C1-32d hinders fear-based HOA formation. g) Using our behavior paradigm, we investigated the neural underpinnings of HOA during remote retrieval using multiple approaches including hippocampal lesions before and after the acquisition, chemogenetics to inhibit the PV-IN of dCA1, cFOS-based Large-scale Brain-wide Connectivity Activation Mapping (LaBCAM), and 2-photon in vivo imaging to follow longitudinal variations in the dendritic spines in retrosplenial cortex. We also tested our behavior paradigm for HOA in aged mice and model mice for Alzheimer’s disease. h) For long term discrimination of context memory, animals are exposed to no-shock context B for about 3minutes in the morning and, 6hours later, same animals are then exposed to context A, where they receive one shock after about 2minutes of initial exposure. Training in discrimination paradigm is conducted for three consecutive days and animals are tested after a month in BA-C or C-BA order across two days. Context B and A are separated by 6hours both during training and testing. i) BA-C subgroup tested in B and A at on day 31, and in C on day 32 after discrimination training show overall significantly different freezing across the contexts (1-w RM ANOVA, F_2,20_ = 40.65, P>F = 8.99E-8, n = 22). Freezing in no-shock context B (20.74±3.02) is significantly higher than C (6.39±1.27) in BA-C order (Post-hoc Tukey test: B vs A: |t|_42_ = 9.11, P > |t|= 2.18E-7; B vs C: |t|_42_ = 5.89, P > |t|= 4.38E-4). Freezing in A (42.94±4.24) is significantly higher than both B and C (Post-hoc Tukey test: A vs C: |t|_42_ = 14.99, P > |t|= 0). j) C-BA subgroup tested in C on day 31 and B-A on day 32 shows significantly higher freezing across contexts (1-w RM ANOVA, F_2,20_ = 21.17, P>F = 1.15E-5, n = 22) although fear in C (18.1±2.47) is similar to fear in B (22.73±2.73), Post-hoc Tukey test: |t|_42_ = 1.97, P > |t|= 0.35; but lower than A(37.17±3.39), Post-hoc Tukey test: |t|_42_ = 8.10, P > |t|= 2.81E-6. k) Context C freezing for C-BA testing order is significantly higher than BA-C (1-w ANOVA, F_1,42_ = 17.76, P>F = 1.30E-4, n = 22 each). l) Context C freezing for C-BA testing order is significantly lower than C1-33d test in ABC (C1-31d(C-BA)-Dis: n = 22, 18.10±2.47; C1-33d(ABC)-Gen: n = 64, 30.99±1.83; 1w-ANOVA: F_(1, 84)_=14.02, P>F = 3.30E-4). Box width represents SEM, error bars represent SD, dashed line inside the box shows the mean and solid line shows the median (% freezing). Grey solid circles represent the fear for individual animals. *** represents p ≤ 0.005** represents p ≤ 0.01. ‘A2-31d’ represents the number of exposures to context A after 31days of training.

Using this rationale, we measure HOA mediated transfer of fear when we test the animal in a context **C**. This context is farther in content similarity from the training context **A** but shares some similarity with the uncertain context **B** where HOA are formed. Since contemporaneous fearful experience resulting from remote retrieval in the training context **A** mediates the conditioning (acting as preconditioned CS), altering the sequence of exposure i.e. retrieval in uncertain context **B** followed by retrieval in original training context **A** should not result in HOA. This is because there is no prior fearful experience that is concurrent, such as previous day’s retrieval in training context, for the animal to associate during remote retrieval in the uncertain context **B**. In other words, we would see HOAs to be formed only when the training context is retrieved first followed by retrieval in a somewhat similar context (**ABC** testing order) and not when the training context is retrieved after the similar context (**BAC**). We also note that this requires the context representation is deconstructed to its elements, thus making the effect more prominent during the remote rather than recent retrievals. Our first set of experiments directly tests the HOA formation and its order dependence.

We train animals in context **A** and subsequently test for remote retrieval in contexts of decreasing similarities starting with the context **A** (original training context) on 31^st^ day, then in context **B** (similar to **A** with partially overlapping features) on 32^nd^ day, followed by context **C** (similar to **B** but distinct from **A**) on 33^rd^ day since training. We tested the remote contextual memory in both **ABC** as well as **BAC** testing order using two distinct groups of animals (Figure 1a, b). Animals exhibit significant freezing during memory retrieval in contexts **A** (n = 64, mean ± SEM; 60.63 ± 2.24) as well as B (42.75 ± 2.07) suggesting that the original fear memory from training is retained long-term (Figure. 1c). However, when tested for context specificity, the animals failed to discriminate the training (**A**) and related context (**B**) only when the related non-training context **B** was tested before **A** (Figure 1d: n = 39, **B**: 50.52 ± 2.90% vs. **A**: 51.02 ± 2.94%; |t|_76_ = 0.25, P > |t| not significant), whereas animals tested in **A** before **B** retained context specificity.

Next, to test if HOA is acquired during sequential exposure to **A** and **B** and if there is a dependence on testing order (**AB** vs **BA**), both groups were subsequently tested in context **C**. Consistent with our premise, we observed a retrieval order dependent increase in freezing for context **C** as it was significantly higher in the **ABC** cohort than in the **BAC** cohort (n = 64, AB**<u>C</u>**: 30.99 ± 1.83 vs. n = 39, BA**<u>C</u>**: 20.24 ± 2.59; F_(1, 101)_ = 12.09, P>F = 7.50E-4). This retrieval order-dependent effect was absent during recent memory tests conducted within 24 hours of training as we find the **C** freezing to be closer to baseline in these recent memory tests (Supplementary Figure 2, 3). Absence of fear in context **C** during recent **ABC** retrieval establishes that the fabrication of fear (via HOA) for a novel context (**C**), requires the context-fear memories to undergo SysCon.

Then, we ruled out the possibility that retrieval order dependent freezing in context **C** could have resulted from generalization or lack of discrimination. If this were to be the case, we would expect the freezing in **C** to be lower for the **ABC** testing order, when animals show discriminability between **A** and **B** (Figure 1c). Instead, we discovered that freezing in context **C** was higher when the memory was tested specifically in the **ABC** order (Figure 1e) suggesting discrimination did not mediate expression of fabricated fear in context **C** (**C** in AB**C** vs BA**C**: F_(1, 101)_ = 12.09, P>F = 7.50E-4). We further tested context **C** freezing after discrimination training, to substantiate that it is the uncertainties associated with remote retrieval that result in HOA (Figure 1h). Previous studies have shown that when trained explicitly to discriminate between related contexts, the animals can discriminate the training and testing contexts remotely with no uncertainty^60^. We took advantage of such training and tested if the fear could be transferred to **C** context in such a scenario following explicit discrimination training between **A** and **B** (Figure 1h). We observed that while the animals can discriminate well between contexts **A** and **B** after explicit discrimination training, just having the ability to discriminate, and thereby eliminating the uncertainty, prevented the increase in fear for context **C** (Figure 1i, j, k, l), when tested after **B** and **A**. It is important to note that low **C**-freezing is not due to discrimination training-based reduced ability of animals to associate fear with a novel context **C**. We observe similar fear recall in contexts **C** and **B** when tested in **C-BA** order after the same discrimination training indicating intact ability of animals for fear transfer involving a novel context **C** (Figure 1j). These observations demonstrate that sequential retrieval in the **ABC** order is necessary but not sufficient for HOA. Thus, we conclude that uncertainty resulting from generalized nature of consolidated memory is necessary for the transfer of fear to neutral context, likely creating false memory percepts. Next, we directly tested the necessity of exposure to context **B** during remote retrieval on HOA based fear transfer.

### Retrieval in an Uncertain context is necessary for HOA

During **ABC**-testing order, we hypothesized HOA develop between the overlapping and non-overlapping elements of contexts **B** and **A**. Since such an association requires both a fearful experience of the recent past and the current uncertain experience, we predict eliminating the uncertain experience should eliminate the transfer of fear via HOA. Consistent with this hypothesis, when we skip the exposure to uncertain context **B** (Figure 1f), as remote fear memory is tested in **A** on day 31 (**A**2-31d: 56.56 ±5.05) and **C** on 32 (C1-32d: 25.62 ±3.22), without the intervening exposure to **B**, the animals show lower freezing that is comparable to routine freezing in **C** tested at 24Hrs (Supplementary Figure 2b).

Further if the remote retrieval experience is resulting in transfer of conditioning to naïve and unique elements then we hypothesize that such a transfer can also happen for safety. Because context **C** is less aversive than **A**, its retrieval immediately after **A** in the **AC** order may instead confer safety properties onto the unique features of **C** while actively preventing subsequent aversive HOA acquisition. Consistent with the safety learning hypothesis^63^, we find that the subsequent exposure to **ABC** order, after **AC** test (i.e. **AC-ABC)**, does not show any appreciable increase in context **C** freezing (Figure 1f, **A**3-33d (44.59±5.64), **B**1-34d (28.89±3), **C**2-35d (20.49±2.62); **C**1-32d vs. **C**2-35d: post hoc Tukey’s test: |t|_48_=1.48, P>|t|=not significant), demonstrating that prior **AC** exposure occludes subsequent fear-based HOA (Supplementary Figure 4).

Overall, low freezing observed in context **C** in the absence of intervening **B** exposure suggests that the nature of the concurrent related experience influences the behavior during the remote retrieval. **C**1-32d freezing without intervening context **B** (Figure 1f) is significantly lower than the HOA dependent increase in **C**1-33d freezing during remote testing in **ABC** order (Figure 1c). Thus, the increased freezing in **C** context seen during remote retrieval in **ABC** requires that the test in uncertain context **B** occurs after **A**. Together, these results establish that HOA requires **B** to be encountered as a state of uncertainty, immediately (within 24-hours) following retrieval of the fearful memory in **A**. The nature of the intervening retrieval experience, whether aversive or safe, determines the direction of associative transfer to novel context **C**. This is consistent with our interpretation of the **BAC** results. During **B** retrieval in the **BAC** order, the overlapping elements of **B** have not yet been associated with a recent fearful experience in **A**, and therefore carry no associative weight to transfer fear to the non-overlapping elements of **C** through HOA ^46,52^.

### Memory uncertainty triggered HOA formation during remote retrieval requires hippocampus

Remote memory of a training event and the associated context is thought to be generalized^11,30^. SysCon renders remote contextual fear memory hippocampus-independent for expression, yet the hippocampus remains essential for encoding new associative information during retrieval^25^. Retrieving the fear training context remotely constitutes an ongoing contemporaneous experience, and such an experience is essential for HOA. Since HOA requires the animal to encode new information during the uncertain retrieval of context **B** soon after a fearful experience in **A**, we reasoned that hippocampal integrity at the time of remote retrieval, not merely at encoding, is the critical variable. An inability to experience fear during remote retrieval in training context should prevent HOA formation. Specifically, since hippocampus is known to be essential for forming memories of life events, an animal unable to engage hippocampal encoding mechanisms during **ABC** retrieval should fail to form HOA, even if the systems-consolidated fear memory itself remains intact.

To test if HOA encoding requires hippocampus, we performed localized Ibotenic acid lesions of the dorsal hippocampus three weeks after initial training to ensure SysCon was complete, followed by one week of recovery before sequential **ABC** testing (Figure. 2a; lesion: n = 12, sham: n = 18, Supplementary Figure 5). Both lesion and sham groups exhibited robust freezing in the training context **A** and the uncertain context **B**, confirming that systems-consolidated remote memory was intact in the absence of hippocampus (Figure. 2b). This is consistent with prior work establishing hippocampus-independence of remote contextual fear expression^21,64^. Hippocampus lesion group tested in **ABC** order (L-**A**2-31d, L-**B**1-32d, L-**C**1-33d) shows overall difference in freezing across contexts (Figure 2b: 1-w RM ANOVA, F_2,10_ = 11.75, P>F = 0.0023, n = 12). However, context discrimination across **A** and **B** is lost in absence of hippocampus (|t|_22_ = 0.84, P>|t| not significant) as animals show similar freezing in both **A** and **B**. Still, animals were able to discriminate the novel context **C** from both **A** (|t|_22_ = 4.57, P>|t| = 0.01) and **B** (|t|_22_ = 5.75, P>|t| = 0.001) showing intact remote memory retrieval in absence of hippocampus. Thus, clearly demonstrating that systems-consolidated memory for the training context was present but that the uncertainty signal carried by **B** was not sufficient to drive HOA in the absence of hippocampus. Sham-lesion group tested in **ABC** order shows discrimination with different freezing across contexts (Figure 2c: 1-w RM ANOVA, F_2,7_ = 20.92, P>F = 0.001, n = 9). Remote fear recall in **A** is significantly higher than **B** (|t|_16_ = 2.72, P>|t| = 0.045) and **C** (|t|_16_ = 5.57, P>|t| = 0.0001), while freezing in **B** is also higher than **C** (|t|_16_ = 2.85, P>|t| = 0.029). However, the freezing in context **C** was significantly higher in sham lesion group compared to the hippocampus lesion group for **ABC** order (Figure 2d: 1w-ANOVA, F_1,19_ = 6.005, P>F = 0.024, t = −2.45). Thus, indicating the HOA mediated confabulated fear memory for context **C** requires intact hippocampus prior to remote retrieval. Animals tested in **BAC** order with sham lesions (vehicle only) confirm the order specificity of HOA formation as they show overall freezing difference (Supplementary Figure 6: 1-w RM ANOVA, F_2,7_ = 12.38, P>F = 0.005, n = 9) with similar freezing between **B** and **A** (|t|_16_ = 0.896, P>|t| not significant) and freezing in **C** still different from both **B** (|t|_16_ = 5.13, P>|t| = 0.0003) and **A** (|t|_16_ = 4.24, P>|t| = 0.0018).

**Figure 2.**
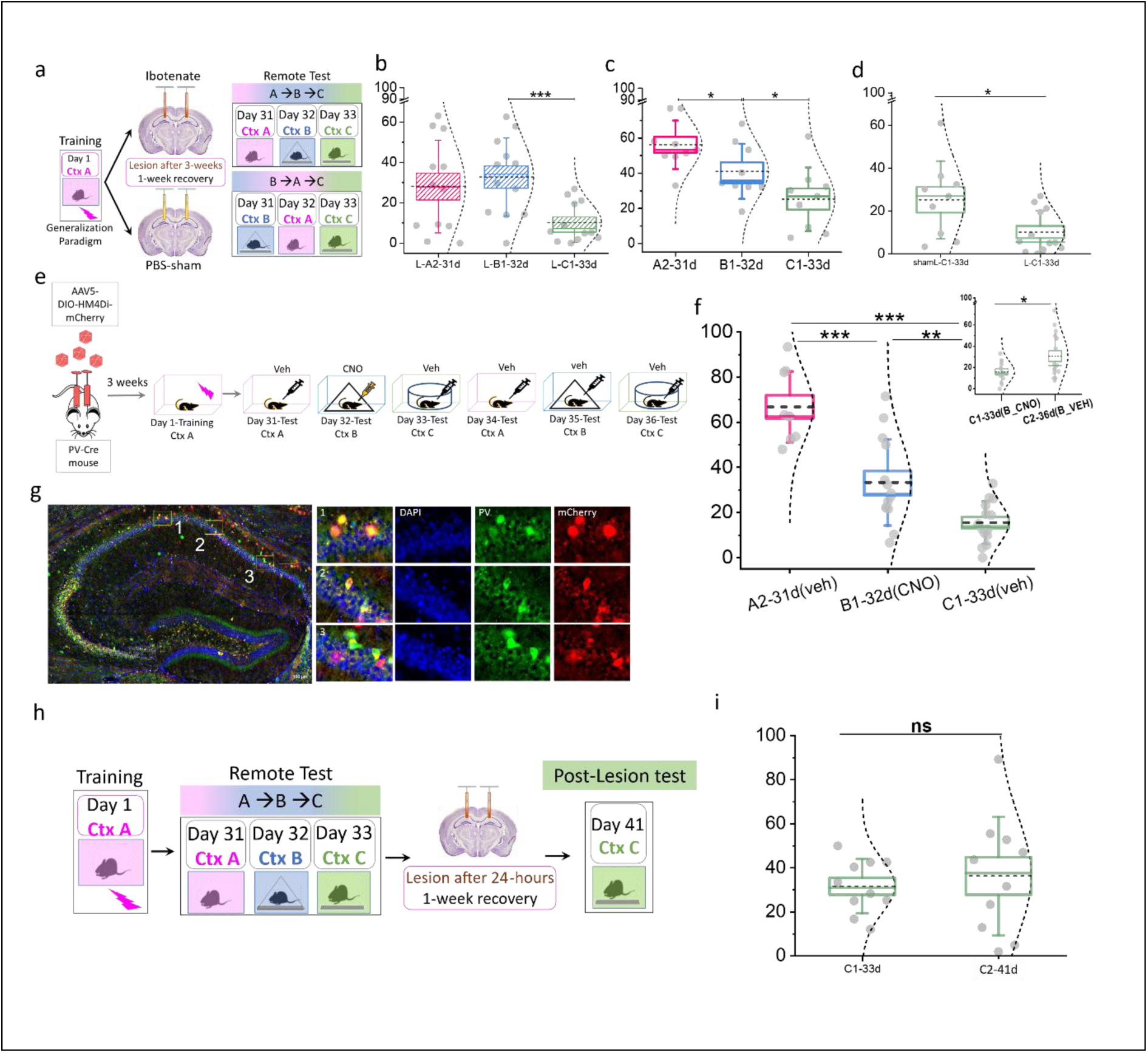
Role of hippocampus in acquiring higher order associations through SOC during remote retrieval. a) Animals (n = 30) are trained in generalization paradigm and after three weeks of systems consolidation, dorsal hippocampus is lesioned in a subgroup of animals (n = 12), while sham-lesions are performed in remaining animals (n = 18, 9 each for sham ABC (1c) and sham BAC (Supplementary Figure 5)). b) Hippocampus lesion group tested in ABC order shows overall difference in freezing across contexts (1-w RM ANOVA, F_2,10_ = 11.75, P>F = 0.0023, n = 12). Testing-order and feature-overlap dependent context discrimination across A and B is lost in absence of hippocampus (|t|_22_ = 0.84, P>|t| not significant) as animals show similar freezing in both A and B. Still, animals were able to discriminate the novel context C from both A (|t|_22_ = 4.57, P>|t| = 0.01) and B (|t|_22_ = 5.75, P>|t| = 0.001) showing intact remote memory retrieval in absence of hippocampus. _c)_ Sham-lesion group tested in ABC order shows testing order-based discrimination with different freezing across contexts (1-w RM ANOVA, F_2,7_ = 20.92, P>F = 0.001, n = 9). Remote fear recall in A is significantly higher than B (|t|_16_ = 2.72, P>|t| = 0.045) and C (|t|_16_ = 5.57, P>|t| = 0.0001), while freezing in B is also higher than C (|t|_16_ = 2.85, P>|t| = 0.029). d) Context C freezing is significantly different for ABC order across sham and lesion group (Two-sample t-test: P>|t|: 0.01206) e) PV-cre mice with AAV-based inhibitory DREADD expression in dCA1 were trained in context A with one shock generalization paradigm. During remote test in ABC order, CNO or vehicle was systemically administered before testing in B, leading to inhibition of PV-Cre interneurons in dorsal hippocampus just before HOA acquisition. f) PV-cre animals with chemogenetic inhibition of dCA1 were trained in A, and then subsequently tested in remote AB_CNO_C and AB_VEH_C order. Animals showed context discrimination with decreasing fear (1-w RM ANOVA, F2,7 = 158.30, P>F =1.48E-6). Freezing in A was considerably higher than B and C, A2-31d (66.77± 5.23, n=9) and B1-32d (33±6.56, n=14), C1-33d (15.56±3.10, n=14). Post hoc Tukey’s test showed the significant difference in freezing percentage across different contexts. Animals showed higher freezing in A2-31(VEH) compared to B1-32d(CNO), (|t|_16_=13.66, P>|t|=6.00E-8) and C1-33d(VEH) (Post hoc Fisher’s test since Tukey’s test value was zero: |t|_16_=21.35, P>|t|=6.90E-11), and B1-32d(CNO) is higher than C1-33d(VEH) (|t|_16_=7.68, P>|t|=1.54E-4). Inset: Freezing between C1-33d(VEH) and C2-36d(VEH) was significantly different (Two sample T-test: |t|_28_=-2.34, P<|t|=0.013), where animals showed higher freezing in C2-36d vs. C1-33d, indicating suppression of PV-IN activity conditionally disrupts the HOA acquisition. g) Coronal dorsal hippocampus section with expression of hM4Di-mcherry (red) in Parvalbumin-interneurons (green). Multiple ROIs in the section were magnified further to show the colocalization and counterstained with DAPI (Blue). h) Animals (n = 12) trained in generalization paradigm are tested for remote retrieval in ABC order and undergo excitotoxic hippocampus lesions after C1-33d test, within 24-hours of the acquisition of SOC-based higher order associations (i.e., 34 days after training). Lesioned animals are allowed ∼a week of recovery from surgery before testing the retention of SOC-based learning during remote retrieval. Two of the animals were excluded due to imperfect lesioning. i) Animals with dHpc lesion ∼24 hrs after HOA acquisition show high freezing for context C (35.46 ± 2.98). Interestingly, C2-41d freezing is similar to the C1-33d test conducted before the lesions suggesting rapid consolidation of higher order associations (|t| = 0.1, P>|t| not significant).

### HOA formation is dependent on CA1 PV interneuron activity

Having established that hippocampal integrity at the time of uncertain retrieval is necessary for HOA, we next asked which hippocampal circuit element gates the uncertainty signal that facilitates HOA formation. Interestingly, CA1 inhibitory interneurons have been shown to play a prominent role in memory formation and affect memory specificity and discrimination without affecting the memory strength^65,66^. Parvalbumin-expressing interneurons (PV-INs), GABAergic inhibitory cells in dorsal CA1 play well-established roles in memory consolidation, network oscillation coordination, and context-specific memory encoding^66–69^. Critically, CA1 PV-IN activity was recently shown to control the precision of contextual memory representations, as reducing the feedforward inhibition shifts CA1 encoding from high-resolution, context-specific representations to low-resolution, generalized representations^70^. We reasoned if PV interneurons can affect memory specificity, inhibiting the PV interneurons would prevent the animal differentiating similar contexts, and thus should remove the uncertainty arising from the second context **B** that resembles closely with the training context **A**. In effect inhibiting PV interneurons should prevent the HOA formation. We hypothesize that the chemogenetic suppression of CA1 PV-INs specifically during **B** exposure, should prevent HOA without disrupting the systems-consolidated fear memory itself. Hence, we probed the necessity of distinguishing the uncertain contexts as being different from the training context for HOA formation using chemogenetics.

To test this, we expressed inhibitory DREADD in PV interneurons of CA1 region of PV-Cre animals (Figure 2e, g, Supplementary Figure 7). Colocalization analysis confirmed that 86% of endogenous PV-INs expressed the DREADD receptor, with minimal off-target expression (Supplementary Figure 8a, b). Animals were trained in context **A** and, after 31 days, tested remotely in context **A** (Figure 2e). On day 32, DREADD activator clozapine-N-oxide (CNO; 3mg/kg, IP) was administered prior to context **B** exposure to selectively inhibit dorsal CA1 PV-INs during the uncertain retrieval window, followed by testing in context **C** on day 33. CNO-treated animals showed significant differences in freezing across the **ABC** sequence, demonstrating context discrimination with decreasing fear (Figure 2f: 1-w RM ANOVA, F_2,7_ = 158.30 P>F =1.48E-6, n = 9). Freezing in **A** (66.77± 5.23, n=9) was significantly higher than **B** (33±6.56, n=14; |t|_16_=13.66, p>|t|=6.00E-8) and **C** (15.56±3.10,n=14) (|t|_16_=21.35, P>|t|=6.9E-11). Freezing in B was also significantly higher than **C** (|t|_16_=7.68, P>|t|=1.54E-4). Importantly, when the same animals served as their own controls with vehicle administration on day 35 prior to **B** exposure, context **C** freezing was significantly elevated relative to the CNO condition (Two sample t-test: |t|_28_=-2.34, P<|t|=0.013) (Supplementary Figure 8c). These findings establish that CA1 PV-IN activity during context **B** retrieval is necessary for HOA formation. PV-INs maintain the representational precision that allows the animal to register the uncertainty between contexts **B** and **A**, the precondition for higher-order associative encoding. Overall, our hippocampal lesion and PV-IN chemogenetic data converge on a two-component necessity for HOA: intact hippocampal engagement during remote retrieval, and PV-IN-dependent precision encoding that registers the uncertainty of context **B**. We next asked whether memories formed through this mechanism undergo SysCon to become enduring, hippocampus-independent traces.

### Uncertainty mediated HOA acquisition shows rapid systems consolidation (SysCon)

Classical SysCon proceeds over weeks to months^11,12,25^, however, the timing of hippocampal disengagement can vary substantially with the nature of the memory being consolidated^23,25,71^. False memories formed through artificial engram manipulation become hippocampus-independent within days of formation^72^, raising the possibility that associatively generated false memories, including those formed through HOA, may consolidate on an accelerated timescale. Consistent with this, RSc lesions which abolish rapid consolidation of standard contextual fear memory^73^ also impair HOA expression, suggesting that the RSc-mediated consolidation mechanism engaged during HOA may be distinct from that of first-order fear memory^62,74^.

To directly test whether HOA-based false memories undergo rapid SysCon, we trained a separate cohort in the **ABC** testing order (Figure 2h, Supplementary Figure 9). Following context **C** testing on day 33: these animals (n = 12) received dorsal hippocampal ibotenic acid lesions 24 hours after **ABC** testing (on day 34, i.e. 34 days post-training). After one week recovery, they underwent context **C** test on day 41 (Figure 2i). Statistical comparison of context **C** freezing across the lesion (**C**2-L-41d) and pre-lesion tests (**C**-33d) revealed no significant difference (|t| = 0.1, P>|t| = not significant). This consolidation timescale appears markedly faster than that observed for standard contextual fear memories, which typically require weeks of hippocampal engagement^11,21^. The rapid hippocampal disengagement of HOA memories is consistent with the pattern separation demands of HOA encoding^23,71^. The higher-order association between context **B** features and the reactivated **A**-US engram requires hippocampal pattern separation during the uncertain retrieval window. The resulting false memory trace is rapidly transferred to RSc-dependent neocortical storage^73,75^, bypassing the extended hippocampal consolidation period required for directly encoded fear memories.

Next, we reasoned if the remote testing in context **C** elicits fear only in **ABC** testing order and not in **BAC**, then the underlying brain wide networks involved should be different despite the external environment being the same (Context **C**). We investigate the brain wide networks involved in remote retrieval triggered HOA using **LaBCAM** (Large-scale Brainwide Correlated Activity Mapping), a lab-developed machine learning based algorithm for detecting correlated activity changes across the brain regions.

### Whole brain mapping and network connectivity

The brain regions recruited during retrieval of HOA-induced fear memory are unknown. We investigated differential activation of these brainwide networks during remote context **C** exposure across the **ABC** (HOA-induced fear) and **BAC** (no HOA-induced fear) testing orders. We used brainwide cFos mapping for identifying co-active networks underlying memory retrieval, as inter-regional correlation of cFos intensity across animals provides an established proxy for identifying putative co-active networks^61,76–81^. We applied LaBCAM to identify brain regions whose cFos activity co-varies across animals during HOA induced fear expression (Figure 3a, Supplementary Figure 10, 11). Animals trained in the generalization paradigm were tested in either the **ABC** or **BAC** retrieval order, perfused 90 minutes after context **C** testing, and immune-stained for cFos (Figure 3a). A random forest classifier implemented in ILASTIK^82^ was used to identify cFos-positive cells, and median cFos intensity was quantified across 20 brain regions spanning cortical, subcortical, and deep brain areas including dorsal hippocampus, thalamic nuclei, and RSc (Figure 3b,c; Supplementary Figure 10; see Methods).

**Figure 3.**
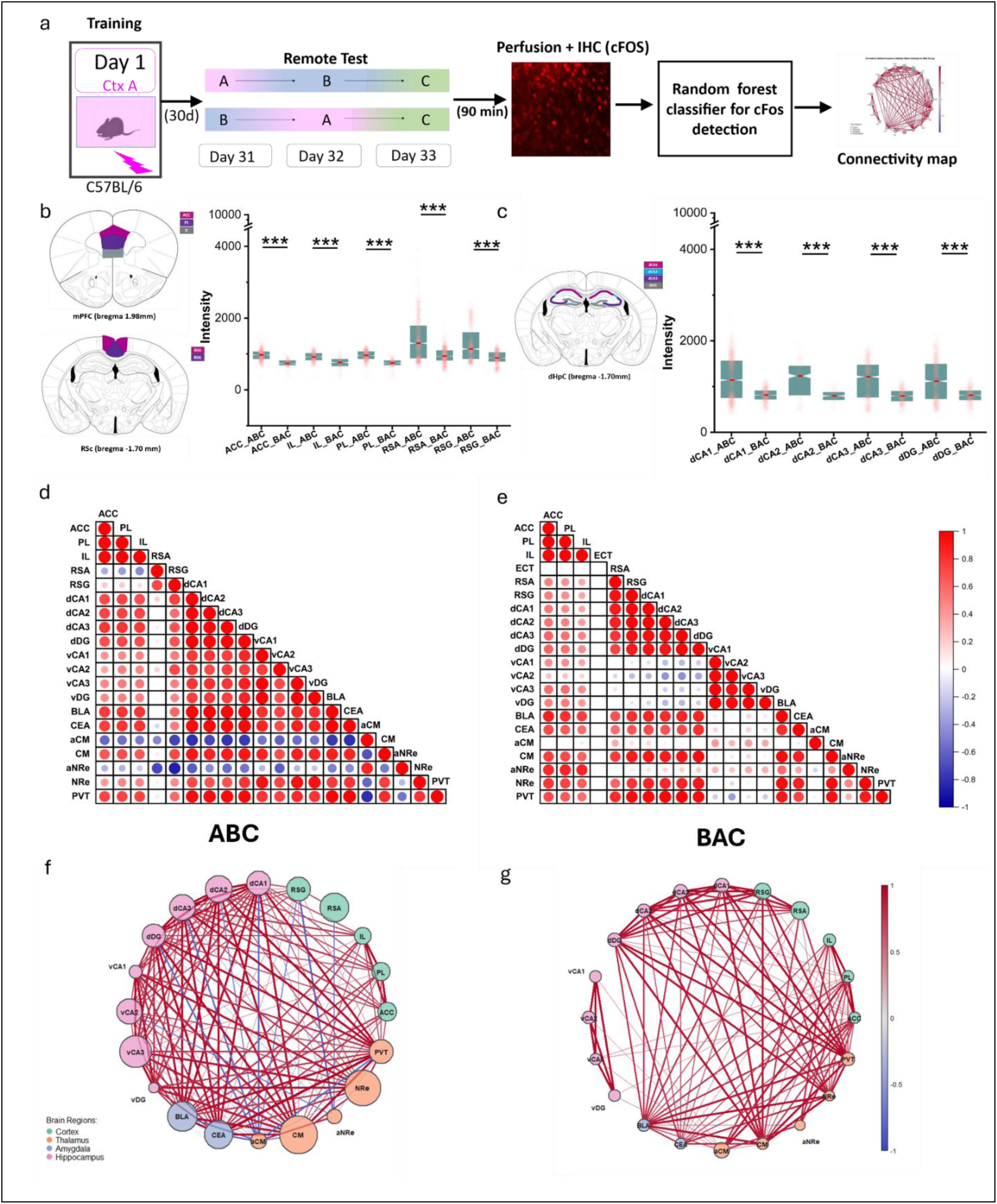
Large-scale Brain-wide Correlated Activity Mapping (LaBCAM) during HOA-based memory retrieval. a) Diagram illustrating the experimental workflow for **LaBCAM**: Post-CtxC testing, animals were perfused, and brain sections from regions like Cortex and Dorsal Hippocampus were immuno-stained for cFos. A random forest classifier was used to detect cFos-positive cells, enabling whole-brain activity mapping and construction of connectivity maps across regions. b) All cortical regions—ACC (Anterior Cingulate Cortex), PL (Prelimbic Cortex), IL (Infralimbic Cortex), RSA (Agranular Retrosplenial Cortex), and RSG (Granular Retrosplenial Cortex)— exhibited significantly higher cFos mean intensity when tested in ABC order (Kruskal-Wallis test with Dunn’s post hoc: ACC: mean rank difference 1368.84, Z=31.68, P=2.32E-22; IL: mean rank difference 677.89, Z=19.57, P=2.61E-85; PL: mean rank difference 1319.52, Z=32.98, P=1.181E-238; RSA: mean rank difference 1735.61, Z=34.69, P=1.07E-263; RSG: mean rank difference 1307.34, Z=30.58, P=1.73E-205). c) Similarly, dorsal Hippocampus subregions (dCA1, dCA2, dCA3, dDG) showed elevated cFos intensity in ABC order compared to BAC order (dCA1: mean rank difference 1217.47, Z=25.38, P=3.74E-142; dCA2: mean rank difference 193.45, Z=11.88, P=1.373E-32; dCA3: mean rank difference 900.10, Z=24.76, P=1.98E-135; dDG: mean rank difference 861.98, Z=21.61, P=1.30E-103), indicating a potential role in HOA-induced memory retrieval. d,e) Pairwise cross-correlation for ABC (left) vs. BAC (right): Colours in the circular areas represent extent of correlations; Pearson coefficients (r). f,g) The connectivity map displays strong correlations (r ≥ +0.5 or r≤-0.5) only, with major brain subregions colour-grouped, node sizes proportional to average cFos intensity, and correlation strength shown by line transparency and thickness of connecting lines.

To gain in-depth insights into potential functional connectivity within these brain networks, inter-regional Pearson correlation matrices were constructed separately for **ABC** and **BAC** groups (Figure 3d, e), where nodes represent median cFos intensity per brain region and edge thickness reflects the magnitude of pairwise correlation across animals. Positively correlated region pairs are shown in red and negatively correlated pairs in blue. We identified a high-degree positive correlation network (p < 0.001) spanning dorsal hippocampus, ventral hippocampus, and amygdala that was selectively present in the **ABC** group and absent in the **BAC** group, suggesting that HOA-induced fear recall in context **C** engages a co-active network overlapping with established fear memory expression circuit (Figure 3c, Supplementary Figure 10b-d). Notably, despite showing markedly elevated cFos activation in **ABC**-order animals, reflected in larger node size, the RSc was significantly less correlated with other regions in this network (Figure 3d-g), distinguishing it from the hippocampal-amygdalar co-activation pattern and suggesting it operates as an independent neocortical node during HOA expression rather than as an integrated component of the distributed fear network^61,75^.

### Retrosplenial cortex (RSc) is necessary for HOA formation

RSc is known to encode and store contextual and spatial information associated with an event^75,83,84^, and to be active during schema-based learning and retrieval^85^. Functional integrity in RSc is essential for the rapid consolidation of fear memories, with macromolecular synthesis inhibition in RSc impairing both recent and remote fear memory expression^62,73^. However, whether RSc is necessary for the formation and retrieval of higher-order associations, where an indirectly conditioned context acquires fear through associative inference rather than direct aversive experience, is untested. Being known for its role in context dependent conflict resolution^86^, and our results indicating higher activity during retrieval of context **C** in **ABC** and not in **BAC** order we wanted to test the necessity of RSc for the retrieval of HOA driven memory in **C** context.

We chemogenetically inhibited excitatory neurons of RSc specifically during context **C** retrieval in the **ABC** sequence (Figure 4a, b). Animals were infused with a Cre-dependent inhibitory DREADD (hM4Di) targeted to RSc excitatory neurons post-training and received either vehicle (Figure 4c) or CNO (Figure 4d) prior to context **C** testing (see Methods). Both vehicle (Veh; n = 25) and CNO (n = 16) groups showed significantly higher freezing in the training context (**A**, day 31, Veh: 56.71±3.85, CNO:50.89 <u>±5.02</u>) than in contexts **B** (day 32, Veh:37.88 <u>± 3.81,</u> CNO:33.28<u>±4.3</u>) and **C** (day 33, Veh: 30.04 <u>± 3.37, CNO: 19. 08± 3.8</u>), confirming intact remote fear memory in both conditions (Veh: F₂,₂₃ = 21.97, P < 0.001; CNO: F₂,₁₄ = 32.64, P = 5.35E-6). Critically, subsequent context **C** freezing was significantly lower in CNO-treated animals than in vehicle controls (Inset, Figure 4d, two-sample t-test, P = 0.02), demonstrating that RSc activity during retrieval is causally required for the expression of HOA-mediated component of fear memory in context **C**.

**Figure 4.**
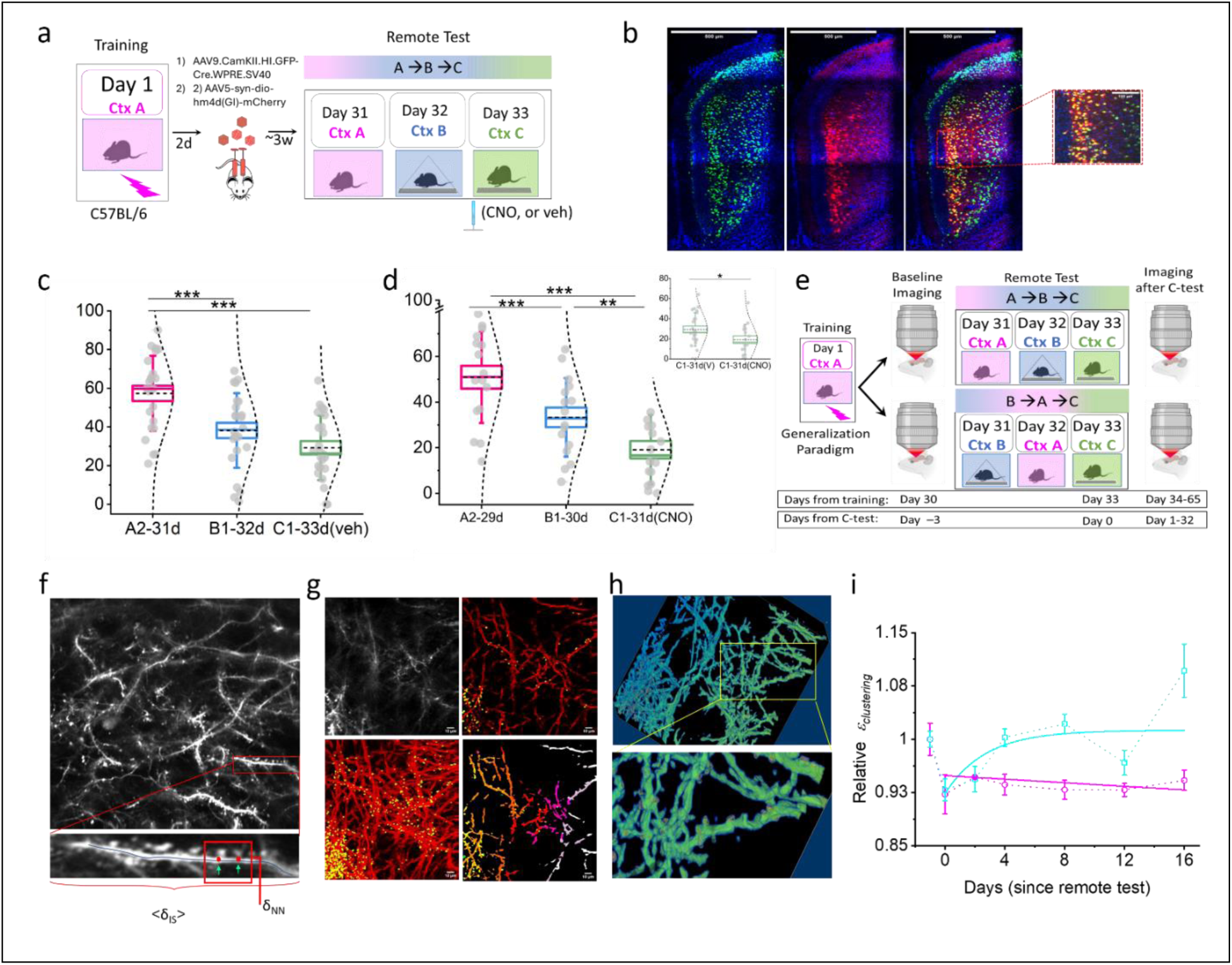
Investigating the role of RSc in HOA acquisition and retrieval using chemogenetic suppression of RSc networks and longitudinal in vivo 2-photon imaging of dendritic spines in RSc. a) Animals (n=41) were trained in three three-shock generalization paradigm and then received the cocktail of viruses (AAV9.CamKII.HI.GFP-Cre.WPRE.SV40 and AAV5-syn-dio-hM4D(Gi)-mCherry) to the RSC to inhibit the excitatory neuronal population. Three weeks post-surgery, animals were tested in remote ABC order and received either vehicle (Veh) or CNO treatment on C1-33d during retrieval of HOA-based fear memory. b) Representative images of AAV9.CamKII.HI.GFP-Cre.WPRE.SV40 (green) and AAV5-hsyn-dio-hm4d(Gi)-mCherry (red) expression in the RSc taken under 20x magnification (0.8 NA), The scale bar is 500 um. A small part of the image was magnified further with 40x magnification to visualize the colocalization. c), d) After training in context A, animals were either tested in ABC_VEH_ or ABC_CNO_ conditions where RSc was chemogenetically inhibited. In both conditions, animals showed contextual discrimination (ABC_VEH_: 1-w RM ANOVA F_2,23_=21.97, P>F= 0.001; ABC_CNO_:1-w RM ANOVA F_2,14_=32.64, P>F=5.35E-6), and exhibited significantly higher freezing during remote retrieval in context A (56.71± 3.85) in comparison to either B (37.58± 3.81) or C (30.04±3.37). However, freezing in contexts B and C was similar for animals that were tested in ABC_VEH_ condition (Post hoc Tukey’s test, mean difference± SEM = 7.53± 4.59, P>|t|=non-significant). In contrast, those tested in ABC_CNO_ showed significant difference in freezing across contexts B (33.28± 4.30) and C (19.08±3.80). (Post hoc Tukey’s test, mean difference± SEM=14.19± 4.56, P>|t|= 0.01). Interestingly, the freezing for context C was significantly lower for ABC_CNO_ condition when RSc was inhibited as compared to ABC_VEH_ condition with intact RSc function (Inset: Two sample t-Test= P>t=0.02), suggesting that RSc suppression hinders the retrieval of HOA induced memory. d) Remote memory test when RSc was suppressed (ABCCNO) just before retrieving the HOA-induced memory. e) Longitudinal in vivo 2photon imaging data reveals that dendritic spines in RSc undergo remodeling post HOA acquisition. RSc of Thy-1 EYFP (+ve) animals was imaged for five different sessions with an interval of four days. f) Three-dimensional segmentation for raw 2-photon images of dendrites and spines was performed using a random-forest classifier-based ML-model in ILASTIK (Supplementary Figure 9). g) Individual dendrites and spines were isolated as unique objects with their spatial coordinates. h) These coordinates were then used to generate a geodesic map. We computed the average interspine distance (<δ_ISD_>) and nearest neighbor distance (δ_NN_) for each spine. As metric of spine reorganization, we computed the normalized difference ((<δISD> - δ_NN_)/<δISD>) to obtain the spine reorganization trends. i) Spine reorganization during ABC remote test (circle, magenta) and BAC remote test (square, cyan) show a persistent reorganization specifically for ABC group after acquisition of higher order acquisitions through HOA. See Supplementary Figure 15 for non-linear fit stats.

The pattern of **B** versus **C** discrimination further supports this interpretation. Vehicle-infused animals (Figure 4c) showed no significant difference in freezing between **B** and **C** (Tukey’s post-hoc: mean difference ± SEM = 7.53 ± 4.59, P > 0.05), consistent with successful HOA-mediated fear transfer to **C**. In contrast, CNO animals (RSc inhibited during context **C** retrieval) (Figure 4d) discriminated clearly between **B** and **C** (14.19 ± 4.56, P = 0.01), with selectively reduced **C** freezing while **B** freezing remained intact, indicating that RSc inhibition specifically abolished the HOA component of fear expression without disrupting retrieval of the directly conditioned memory. We interpret these results as an indication of RSc’s role in retrieval and rapid SysCon of the HOA mediated memory.

### Dendritic reorganization underlying the development of HOA

Having established that RSc activity is causally required for HOA retrieval, we next sought the sub-neuronal structural correlate of HOA memory storage within RSc (Figure 4e). Clustered plasticity, the coordinated reorganization of dendritic spines within discrete dendritic branches, has emerged as a proposed mechanism for efficient information storage in cortical memory circuits, including RSc^83,87–89^. We therefore used longitudinal in vivo two-photon imaging of dendritic spines in Thy1-EYFP mice to probe and follow the changes in dendritic spines within RSc that could facilitate the consolidation of HOA-mediated false memories. We track structural plasticity in RSc across a six-session imaging protocol spanning four weeks following remote test in context **C** on day 33 of **ABC** or **BAC** retrieval protocol (Figure 4e, Supplementary Figure 12, 13; Methods). Baseline imaging was performed before remote testing for both groups, providing within-animal reference for all subsequent comparisons. Geodesic maps of dendrites and spines were obtained (Figure 4f-h, Supplementary Figure 14) with raw 2-photon images using machine learning algorithm-based segmentation (ILASTIK)^82^.

Overall spine density changed only marginally following remote context **C** retrieval in both **ABC** and **BAC** groups. However, inter-spine distances shifted substantially in the **ABC** group, revealing reorganization of the existing spine population rather than net addition. We observe persistent changes in inter-spine distances even with these marginal changes in spine density, suggesting that HOA induced spine clustering in RSc (Figure 4i). We reason that if the spines are undergoing reorganization, then measuring the nearest neighbor distance and comparing it with average expected interspine distance would provide a good measure of spine clustering. To quantify this, we computed δ_NN,_ the average nearest-neighbor distance between spines on a given dendrite, relative to ⟨δ_ISD_⟩, the average inter-spine distance given that dendrite’s spine density. The magnitude of the quantity |(δ_NN_ − ⟨δ_ISD_⟩)/⟨δ_ISD_⟩| provides a density-normalized clustering index, i.e. relative extent of clustering (Relative ɛ_clustering_), a measure of how closer the dendritic spines are given their density (see Methods).

While both the groups show initial decrease in this fraction (Relative ɛ_clustering_), the **BAC** group rebounds to its initial spine clustering state while that of the **ABC** stays persistently declustered. This is consistent with the role of RSc being a context dependent conflict resolver (Figure 4i). We interpret that in **ABC** group, HOA-mediated memory confabulation occurs following a severe conflict between overlapping and non-overlapping features of **B and A**. Encoding and subsequent consolidation of HOA-mediated false memory results in persistent decline in Relative ɛ_clustering_. Whereas, in **BAC** group, RSc solves a moderate conflict to separate context **C** from **B** and **A** during remote tests, reflected in transient decline in Relative ɛ_clustering,_ that reverts back to baseline clustering state soon after. Essentially, we show selective strengthening of projections to RSc during HOA-mediated memory confabulation. Our findings are consistent with the observation of clustering and de-clustering shown as a neural correlate for storing related memories^90,91^, with clustered spines formed during associative learning in RSc selectively stabilized over non-clustered spines at remote timepoints^87^.

### Early-life HOA deficits in APP-PS1 mice model of AD

HOA utilizes the existing memory to form new memory. Thus, any test of HOA would test not only the ability to form new memory but also the ability to retrieve and utilize the prior memory. We reasoned that in a condition where memory consolidation begins to fail gradually, HOA would be disrupted before standard remote memory measures, because any subtle degradation of the existing memory trace would compromise the quality of the retrieval event that initiates HOA. Hence, HOA could serve as a sensitive early indicator in a developing memory disorder. Alzheimer’s disease, characterized by progressive amyloid accumulation and early hippocampal dysfunction that precedes overt memory loss^92,93^, presents precisely this scenario. We therefore tested HOA in APP/PS1 mice and wild-type (WT) littermates at an age when the training context memory remains measurably intact, to determine whether HOA fails before standard contextual fear memory loss due to dementia. We hypothesize that HOA could be affected in these disease models much before the manifestation of memory loss/dementia.

APP/PS1 (AD+ve) and WT (AD-ve) mice were trained in the generalization paradigm and tested remotely in either **ABC** or **BAC** order (Figure 5a, b). AD+ve animals showed intact contextual discrimination in both **ABC** (n=9; 1-way ANOVA, F₂,₇ = 24.11, P = 7.24E-4) and **BAC** (n=12; F₂,₁₀ = 11.44, P = 2.60E-3) testing orders, confirming that systems-consolidated remote fear memory remained measurably intact. In the **BAC** order, freezing in contexts **B** (42.48±7.45) and **A** (46.09 ± 5.58) did not differ significantly in AD+ve animals (Tukey’s post-hoc: P = 0.80), consistent with the retrieval generalization that characterizes remote testing in both genotypes. Critically, context **C** freezing in AD+ve animals did not differ across **ABC** (22.79±4.95) and **BAC** (22.60±3.5) orders (Figure 5a, Inset; two-sample t-test: P = 0.48), **C** freezing was equivalently elevated regardless of retrieval sequence, demonstrating a loss of the sequential order-specificity that normally restricts HOA to the **ABC** order.

**Figure 5.**
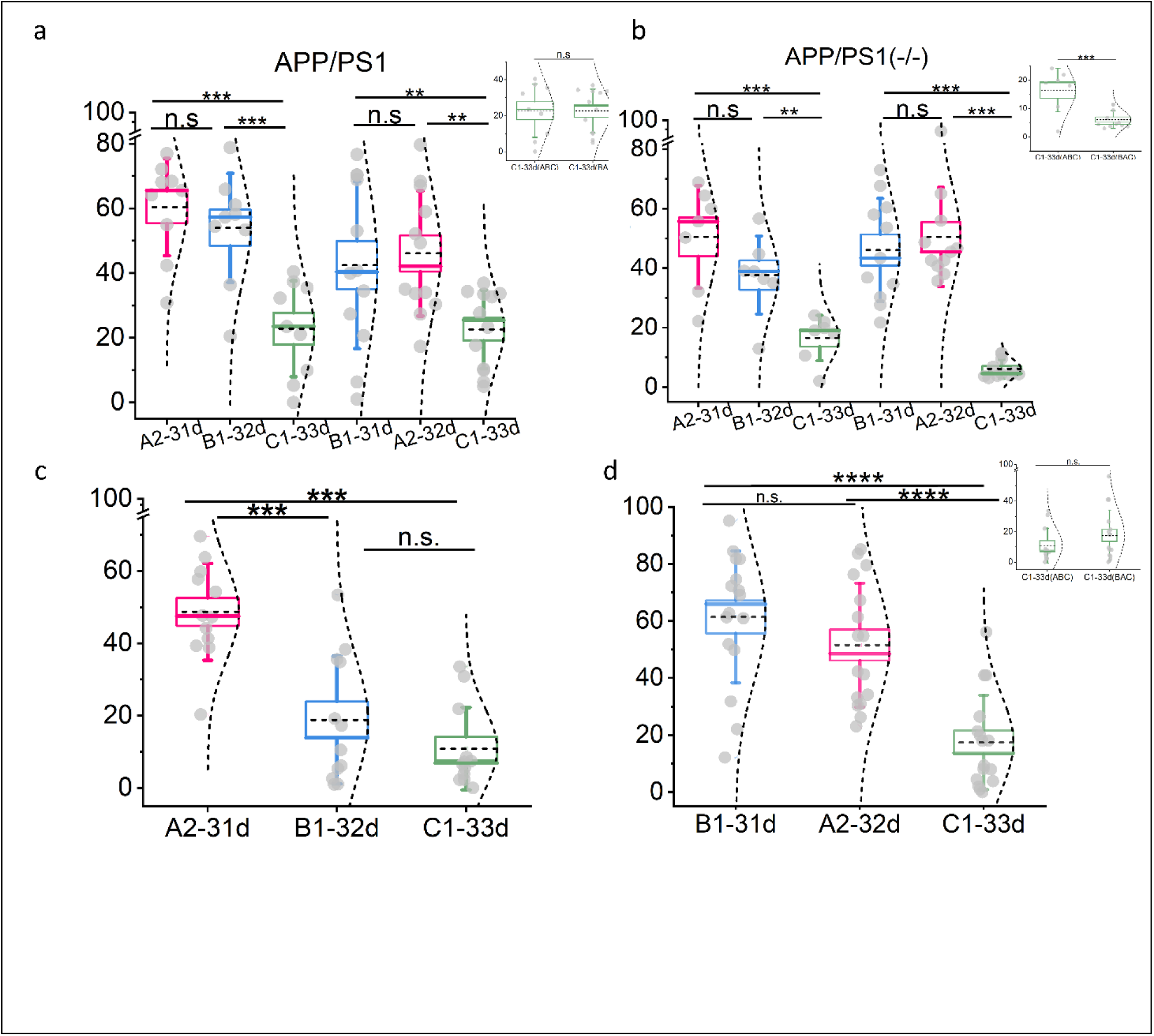
HOA is selectively disrupted in APP/PS1 mice and aged animals through distinct mechanisms. **a)** APP/PS1 (AD+ve) and WT (AD-ve) littermates tested remotely in ABC and BAC order. AD+ve mice show intact contextual discrimination in both ABC (1-way ANOVA, F₂,₇ = 24.11, P = 7.24E-4) and BAC (F₂,₁₀ = 11.44, P = 2.60E-3) orders, with no significant difference in B and A freezing during BAC testing (Tukey’s: P = 0.80), confirming intact remote fear memory. Inset: Context C freezing does not differ between ABC and BAC orders in AD+ve animals (two-sample t-test: P = 0.48, n.s.), demonstrating loss of HOA order-specificity. **b)** In contrast, WT APP/PS1(−/−) littermates show significantly higher context C freezing in ABC than BAC order (F₂,₅ = 59.80, P = 3.22E-4; F₂,₉ = 37.58, P = 4.27E-5; two-sample t-test: P = 4.80E-4), confirming intact order-specific HOA. **Inset:** Context C freezing across ABC and BAC orders for AD -ve animals shows significant difference. **c)** 16–17-month-old C57BL/6 aged mice (n = 28) tested remotely in ABC (n = 12) or BAC (n = 16) order. ABC-tested aged mice show intact remote fear memory and context discrimination, with freezing in A (48.69 ± 3.86) significantly exceeding B (18.77 ± 5.09; 1-way RM ANOVA, F₂,₁₀ = 20.64, P = 2.82E-4; Tukey’s: |t|₂₂ = 7.33, P = 9.70E-5). **d)** BAC-tested aged mice show contextual generalization between B (61.42 ± 5.77) and A (51.53 ± 5.42; 1-way RM ANOVA, F₂,₁₄ = 45.46, P = 7.53E-7; Tukey’s: |t|₃₀ = 3.00, P > 0.05). **Inset:** Context C freezing does not differ between ABC and BAC groups in aged animals (|t|₂₆ = −1.17, P > 0.05, n.s.), indicating global failure of HOA formation regardless of retrieval order, mechanistically distinct from the APP/PS1 phenotype where C freezing is equivalently elevated in both orders. Data shown as mean ± SEM. n.s., not significant.

On the contrary, their APP/PS1(−/−) WT littermates showed robust contextual discrimination in both **ABC** (n=7) (F₂,₅ = 59.80, P = 3.22E-4) and **BAC** (n=11) (F₂,₉ = 37.58, P = 4.27E-5) orders, and showed significantly higher context **C** freezing in **ABC** (16.45±2.89) than **BAC** (6.11±0.95) order (Figure 5b, Inset; two-sample t-test: P = 4.80E-4), substantiating the order-specificity of HOA in normal animals and confirming that this specificity is selectively abolished by APP/PS1 pathology.

The loss of retrieval order-specificity but intact contextual fear memory mirrors the behavioral consequence of CA1 PV-IN inhibition. This is consistent with early amyloid-driven interneuron hyperexcitability disrupting the temporal precision of sequential associative encoding without abolishing the consolidated fear memory itself ^94^. This deficit is detectable while standard remote memory measures remain intact. Hence, the HOA order-specificity paradigm represents a sensitive early behavioral marker for preclinical AD-associated circuit dysfunction, identifiable at a disease stage when conventional contextual fear assays would not show impairment^92,93^.

### Advancing age produces a distinct HOA failure: impaired formation rather than lost order-specificity

We next asked whether normal aging produces a similar or qualitatively different HOA deficit. If aging simply recapitulates the AD phenotype, HOA order-specificity loss would be a non-specific marker of hippocampal decline. However, if aging produces a different pattern, it would suggest that the two conditions disrupt distinct components of the HOA circuit.

HOA-mediated memory encoding requires a previously formed schema^26,27,84^. Though challenging, researchers often use aged mice to study cognitive decline in humans. Aged humans show deficits in episodic specificity and increased reliance on gist-based retrieval that impairs their ability to form new associations linked to established remote memories^95,96^. In aged rodents, impairments in spatial learning, working memory, cognitive flexibility, and novel object recognition are well documented, accompanied by declining hippocampal LTP, dysregulated protein synthesis^97^, and altered synaptic plasticity across hippocampus, prefrontal cortex, and amygdala^98–100^. Whether these age-related changes specifically impair HOA as distinct from generalized memory decline remains untested.

We trained 16-17-month-old C57BL/6 mice (n = 28) in the generalization paradigm and tested them remotely in either **ABC** (n = 12) or **BAC** (n = 16) order. In the **ABC** group, aged mice showed clear contextual discrimination (Figure 5c). Freezing in the training context **A** (48.69 ± 3.86) significantly exceeded that in the related context **B** (18.77 ± 5.09; 1-way RM ANOVA: F₂,₁₀ = 20.64, P = 2.82 E-4; Tukey’s post-hoc: |t|₂₂ = 7.33, P = 9.70E-5), demonstrating that remote fear memory and context discrimination remain largely intact at this age. In contrast, for the **BAC** group (Figure 5d), aged mice showed contextual generalization between **B** (61.42 ± 5.77) and **A** (51.53 ± 5.42) at remote retrieval (1-way RM ANOVA, F₂,₁₄ = 45.46, P = 7.53E-7; Tukey’s post-hoc: |t|₃₀ = 3.00, P > 0.05), consistent with the time-dependent shift from context-specific to generalized remote memory representations documented in both rodents and humans^30,58^.

Critically, context **C** freezing did not differ between **ABC** and **BAC** groups in aged animals (Figure 5d, Inset; one-tailed two-sample t-test: |t|₂₆ = −1.17, P > 0.05). In young animals, **ABC** testing reliably produces elevated **C** freezing relative to **BAC** through HOA-mediated fear transfer; in aged animals, no such order-dependent elevation is detected in either direction.

This pattern is mechanistically distinct from the APP/PS1 phenotype. AD+ve mice lose order-specificity but show equivalent high-**C** freezing in both orders, indicating that HOA-like fear transfer occurs non-selectively. However, aged animals show a global absence of **C** freezing elevation regardless of retrieval order, indicating that HOA formation itself fails. For future investigations, we propose that aging disrupts the hippocampal encoding machinery, specifically the LTP and protein synthesis capacity required during the uncertain **B** retrieval window^98^, such that the CA1 circuitry cannot form the new higher-order association even when the original fear memory is retrievable and the correct sequential structure is presented. This contrasts with early amyloid pathology, which degrades the temporal precision of sequential gating while leaving the associative machinery sufficiently intact for indiscriminate HOA-like transfer. The two translational models thus reveal a double dissociation, AD disrupts *when* HOA occurs, whereas aging disrupts *whether* HOA occurs or not.

## Discussion

The findings presented here establish that the brain’s normal mechanisms for higher-order associative learning can, under precisely defined conditions of remote retrieval and contextual uncertainty, generate a false memory of an event that never occurred. We show that when a systems-consolidated fear memory is retrieved in a context that only partially overlaps with the original training environment, such retrieval provides the uncertainty necessary for second-order conditioning (SOC). The unique features of that uncertain context acquire conditioned fear through higher-order association (HOA), and this false fear memory undergoes rapid SysCon to become hippocampus-independent within 24 hours. The neural architecture supporting this process includes, but is not limited to, CA1 PV interneuron-dependent precision encoding, RSc-dependent retrieval and storage, and a coordinated brain-wide correlated activity pattern restricted to the retrieval order **ABC** that selectively yields HOA-mediated memory confabulation.

### HOA as a mechanism for false memory: ethological versus artificial induction

HOA is an evolutionary mechanism for expanding the causal link between stimuli and secondary reinforcers, and serves a vital role in enhancing the rate of survival. In real-world scenario, learning and hence the survival capacity would be severely limited if it were to depend only on primary reinforcers; HOA enables other agnostic stimuli to acquire the valence. HOA helps the animal to optimize the cost-benefit ratio associated with every exposure to a dangerous or beneficial unconditioned stimulus (e.g., predator, food, or mate). In this study we show that this ability can result in confabulation and trigger false memory formation for an apparent life event that never occurred. Prior work has demonstrated that false memories can be implanted in mice by directly activating a context engram with optogenetics during fear conditioning in a different context, producing conditioned fear in an environment where shock was never delivered^72^. Our findings extend this framework in a critical direction: we show that false memories emerge spontaneously during the normal sequence of remote memory retrieval, without any artificial engram manipulation. However, the mechanism of a false memory formed through artificial, invasive and direct stimulation of neurons in the brain could be different from a memory that is acquired and expressed in near ethological laboratory settings. In our study we show that an experience of having retrieved an old fearful memory in an uncertain context triggers such a conditioning for a new, hitherto unexperienced, context. This bears striking similarity to real world scenario of suspect identification where the victim/eyewitness is asked to identify the perpetrator from a line of possible suspects and illustrates the effect of HOA in forming false memories. HOA-mediated false memory formation, as we show, is gated by the temporal structure of retrieval events, requires hippocampal encoding machinery at the time of uncertain re-exposure, depends on PV interneuron-mediated representational precision, and is expressed through a neocortical storage mechanism involving RSc synaptic reorganization. Our study provides the neural underpinnings of a socio-politically relevant process that underlies real-world memory confabulation^1,3,101^.

### Circuit architecture of HOA

Our results identify a hierarchical circuit for HOA formation and consolidation. At the encoding stage, CA1 PV interneurons are causally required as chemogenetic inhibition of PV-INs during context B retrieval abolishes HOA without disrupting the consolidated fear memory itself^67,68^. This result suggests PV-IN-mediated pattern separation as the cellular gating mechanism for HOA. The unique features of B acquire conditioned fear through second-order association, likely via PV-IN mediated distinction across the contexts during remote retrieval. Moreover, when this discrimination is pharmacologically prevented, the uncertain retrieval event loses its ability to drive HOA.

At the network level, LaBCAM reveals that HOA-inducing retrieval selectively recruits coordinated activity across dorsal hippocampus, ventral hippocampus, and amygdala, a pattern absent in the BAC order, identifying a fear circuit co-activation signature specific to the HOA condition rather than to fear expression per se^76–78^. The RSc, despite being necessary during HOA retrieval, shows markedly lower co-variance with this network. RSc likely operates as an independent neocortical node rather than an integrated component of the distributed fear circuit. Such RSc activity would be consistent with its role as a consolidation relay for rapidly forming associative memories^62,73,75^. RSc chemogenetic inhibition and longitudinal spine imaging together demonstrate that RSc is both causally required for HOA retrieval and undergoes persistent dendritic spine clustering that endures for weeks following HOA acquisition, suggesting the synaptic-level structural substrate of the confabulated memory trace^87,89^.

### Rapid systems consolidation of false memories

Standard contextual fear memories require 3-4 weeks of hippocampal engagement before becoming hippocampus-independent; HOA-generated false memories cross this threshold within ∼24 hours of formation. This accelerated trajectory is consistent with prior observations that artificially induced false memories consolidate rapidly^72^ and that optogenetic reactivation of RSc engram ensembles can replicate SysCon in a single session^75^. We propose that RSc, primed by the HOA encoding event, rapidly consolidates the higher-order association through dynamic modulation of dendritic spine clustering that takes place during the post-retrieval period, eventually, making the false memory resistant to hippocampal interference. This timeline has direct consequences for eyewitness testimony: our data provides a mechanistic explanation for why witnesses who express initial uncertainty at lineup routinely testify with absolute confidence in court. Within 24 hours of the first lineup, a retrieval event in a partially overlapping context, the false identification has consolidated into a hippocampus-independent trace indistinguishable from a true memory ^2,7,8^.

### Translational implications: Alzheimer’s disease, aging, and eyewitness confabulation

Our APP/PS1 data reveal that early amyloid pathology selectively disrupts the sequential gating of HOA, producing C-context freezing in both ABC and BAC orders while leaving direct fear memory intact. Whereas normal aging produces a qualitatively different failure, abolishing HOA formation in both orders without disrupting the underlying fear memory. AD degrades temporal precision while leaving associative capacity partially intact; aging degrades the associative encoding machinery itself, consistent with age-related LTP and protein synthesis failure in hippocampus and prefrontal cortex^98^. The AD phenotype echoes clinical observations of increased false recognition and confabulation susceptibility in early amyloid pathology before overt dementia^92,93^. Mechanistically, PV-IN hyperexcitability in RSc likely disrupts remote fear memory precision in AD mouse models^94^. The HOA order-specificity assay thus offers a behaviorally accessible window into early amyloid-driven circuit dysfunction at a disease stage when standard fear conditioning would not detect impairment.

Beyond neurodegeneration, HOA-based false memory formation is likely operative across domains where associative inference generates beliefs that diverge from direct encoding. For example, HOA could be implicated in post-traumatic stress disorder, in which salient prior trauma primes subsequent uncertain contexts for aversive association; agoraphobia, in which the sequential retrieval structure of panic progressively generalizes to new environments; brand schema formation and consumer decision-making, in which repeated exposure to products in partially overlapping contexts drives affective transfer through HOA rather than direct experience. We have identified and mechanistically dissected a form of false memory that emerges from the brain’s own machinery for higher-order associative learning during remote retrieval. HOA-based false memories are formed through PV-IN-dependent precision encoding, consolidated rapidly through RSc-mediated synaptic reorganization, and retrieved via a coordinated brain-wide, correlated activity pattern restricted to the sequential context structure that generated them. Their failure in distinct ways in APP/PS1 mice and in normal aging defines two clinically relevant modes of HOA circuit disruption. And their parallel to the sequential structure of eyewitness identification procedures positions the HOA framework as a mechanistic account of one of the most consequential sources of false belief in the legal system. The capacity for higher-order associative learning is indispensable for adaptive behavior; its misapplication, as characterized here, is the neural substrate of memory’s most dramatic failures.

## Author contributions

**AS** and **BJ** conceptualized the study. **AS**, **ND**, and **BJ** designed the experiments, performed data analyses, and wrote the manuscript. **AS**, **ND**, **SR**, **TS**, **PA**, **AA**, and **SKu** performed behavioral experiments and transcardial perfusions. **SKu** and **AS** contributed to hippocampal lesion experiments. **ND**, **PA**, and **AA** performed cFos immunostaining, RSc chemogenetic inhibition, and LaBCAM experiments. **ND**, **PA**, **AB**, and **BJ** developed and optimized the LaBCAM analysis. **AS** performed longitudinal two-photon in vivo imaging of RSc dendritic spines with behavior; **SKa** ran the ILASTIK-based analysis pipeline for spine clustering quantification. All authors reviewed and approved the final manuscript.

## Acknowledgement and Funding

We are thankful to Mr. Aritra Dutta for providing initial help with the classifier training, Dr Vikram Pal Singh for contributing to dendrite tracing with simple neurite tracer, Drs Suraj Kumar, Saumitra Yadav, and Vijay R. Singh for supporting multi-photon microscope set-up, and Ashish Das for supporting electronics for custom fear conditioning set-up. We are also thankful to the funding agencies: This work was supported by funding given to **BJ** from SERB (EMR/2017/004155), (CRG/2022/003148), Council for Scientific and Industrial Research grants to **AS** (CSIR-09/079(2590)/2012-EMR-I) and **ND** (CSIR-09/079(2801)/2019-EMR-I), and was carried out in behavior facility supported by DST FIST grant (SP/FST/LS-II/2019/460) awarded to Center for Neuroscience.

