## Supplementary Figures and Tables for "Retrieval uncertainty drives rapid consolidation of confabulated memories in retrosplenial cortex"

### Supplementary Information (Singh-Das et. al. 2026)

### Methods

#### Statistics

All statistical analyses were performed using OriginPro software (OriginLab ver. 9.0 and 10.0). Repeated-measures data were analyzed using one-way repeated-measures analysis of variance (1-way RM ANOVA) followed by post-hoc Tukey's honest significant difference test for pairwise comparisons. Between-group comparisons of context C freezing were performed using two-sample t-tests (two-tailed unless otherwise stated). A significance threshold of $\alpha = 0.05$ was applied throughout. All data are reported as mean ± SEM. Sample sizes (n) refer to individual animals. Statistical test results are reported as F-statistics (ANOVA) or $|t|$ values (t-tests) with degrees of freedom as subscripts. For LaBCAM connectivity analyses, we used non parametric Kruskal-Wallis test with Dunn's post hoc comparison.

#### Animals

Male C57BL/6J wild-type mice (JAX 000664, 2-4 months old) were used for behavior, hippocampal lesion, LaBCAM, RSc inhibition chemogenetics, and contextual fear conditioning experiments. Thy1-EYFP mice (B6.Cg-Tg(Thy1-YFP)HJrs/J, JAX 003782, 8–10 weeks old) were used for longitudinal *in vivo* two-photon imaging of dendritic spines. PV-T2A-Cre mice (Pvalb-T2A-Cre-D, JAX 012358) were used for CA1 PV-IN chemogenetic experiments; animals of both sexes aged 2-6 months were used. APP/PS1 double-transgenic mice (JAX 034829, C3H genetic background; expressing Mo/HuAPP695swe and PS1-dE9) and their wild-type littermates were used as the Alzheimer's disease model; both male and female animals aged 2–5 months were used. Aged cohort experiments used 16-17-month-old male C57BL/6J mice.

All animals were co-housed (2–3 per cage) under a 12:12 hour light:dark cycle with ad libitum access to food and water. Animals were transported to the holding room of the Centre for Neuroscience building for all behavioral experiments. PV-T2A-Cre mice were initially obtained from Jackson Laboratory and subsequently bred in-house at the Central Animal Facility (CAF),

Indian Institute of Science. All procedures were conducted in accordance with the animal ethics protocol approved by the Institutional Animal Ethics Committee, Indian Institute of Science.

#### Context Design

Three contexts (A, B, C) were designed with decreasing featural overlap in the order A > B > C, differing in olfactory, visual, and tactile features. All contexts were housed within sound-attenuated wooden enclosures in the same experimental room, and the same Perspex chamber (inner dimensions: 35 cm × 24 cm × 22 cm, with transparent top, back, and door panels and black Perspex side walls) was reconfigured for each context. A miniature fan was attached to each enclosure to provide constant background noise. Details of context feature compositions are shown in Supplementary Figure 1.

##### *Context A*

White LED overhead illumination. Stainless steel grid floor (32 rods, 2 mm diameter, 10 mm spacing). Odor: 15% acetone solution. White screen covering camera mount on back wall.

##### *Context B*

White LED overhead illumination. Steel grid covered with a laminated white sheet (flat, slippery surface). Triangular polycarbonate insert added as spatial cue, partially attenuating illumination. Odor: 70% ethanol solution.

##### *Context C*

Infrared illumination only (visible LEDs off). Rough-textured surface placed over the flat floor of context B. Triangular polycarbonate insert removed to allow contact with chamber walls. Odor: 10% ethyl acetate solution.

##### *Context C'*

Infrared illumination only (visible LEDs off). Rough-textured surface placed over the flat floor of context B. Triangular polycarbonate insert replaced with open top and open back cylindrical insert that marginally reduced the total area available to explore. Odor: 10% ethyl acetate solution.

#### Behavior Paradigms

#### Handling and Habituation

Each animal was handled for 10 minutes daily for 3–5 days in the experimental room. Animals were not exposed to any training or testing contexts during handling. All training and testing sessions were conducted between 0800 and 1000 hours by the same experimenter, who was blinded during video analysis.

#### Generalization Paradigm

##### *Training*

Animals were individually placed in context A and allowed to explore for approximately 120 seconds. Exploration was followed by three foot-shocks (0.7 mA, 2 seconds each) delivered at 30-second intervals through the steel grid floor, followed by 30 seconds of post-shock exploration. Animals were returned to their home cages after approximately 210 seconds total in context A (Figure 1b, Supplementary Figure 1).

##### *Remote Testing*

Remote memory tests were conducted beginning on day 31 post-training. For ABC testing order, animals were tested in context A on day 31, context B on day 32, and context C on day 33. For BAC testing order, animals were tested in context B on day 31, context A on day 32, and context C on day 33. During each testing session, animals were allowed to explore undisturbed for 120–150 seconds. A subgroup of animals was tested at a recent time point (24–48 hours post-training) in the same ABC or BAC order for comparison with remote memory tests. (Figure 1b, Supplementary Figure 1)

#### Discrimination Paradigm

##### *Training*

Discrimination training was conducted over three consecutive days (Figure 1h, Supplementary Figure 1). Each morning, animals were individually placed in context B (no-shock context) for approximately 200 seconds. Six hours later, the same animals received one foot shock (0.7 mA, 2 seconds) after approximately 135 seconds of exploration in context A (shock context), followed by 30 seconds post-shock before returning to home cage.

##### *Testing*

For remote discrimination testing (day 31–32), animals were assigned to BA-C or C-BA testing order subgroups. In the BA-C order, context B was tested in the morning of day 31, context A in the afternoon of day 31, and context C on day 32. In the C-BA order, context C was tested on day 31, context B in the morning of day 32, and context A in the afternoon of day 32. Recent discrimination tests were performed on days 4–5 post-training using the same structure. (Figure 1h, Supplementary Figure 1)

#### Hippocampal Lesions

##### Pre-Acquisition Lesions

Bilateral dorsal hippocampal lesions were performed by infusion of Ibotenic acid (10 mg/ml in 1× PBS, pH 7.4; 200 nl per site) at three pairs of coordinates: Site 1: AP −1.3, ML ±1.2, DV −2.0 mm; Site 2: AP −2.5, ML ±2.5, DV −2.2 mm; Site 3: AP −3.3, ML ±3.1, DV −4.1 mm (all relative to bregma). Sham controls received equivalent volumes of sterile 1× PBS. Lesions were performed three weeks after training to allow complete systems consolidation. Animals recovered for one week before remote testing in ABC or BAC order (Figure 2a, Supplementary Figure 5).

##### Post-Acquisition Lesions

To test the systems consolidation timeline of HOA-based memories, hippocampal lesions were performed after ABC remote testing on day 33. The remote lesion (RL) group received Ibotenic acid lesions 24 hours after context C testing (day 34) and was subsequently tested in context C on day 41. Animals recovered for one week following lesion surgery before behavior testing. Sham-lesioned animals received 1× PBS infusions on the same schedule. (Figure 2h, Supplementary Figure 5a-c,e)

##### Lesion Surgery Protocol

Animals were anesthetized with isoflurane (5% induction, 2% maintenance). Carprofen (150 µl, 1 mg/ml, subcutaneous) was administered for analgesia. Eye ointment (Neosporin) was applied. After scalp incision and connective tissue removal, infusion sites were drilled using a pneumatic dental drill. A 32-gauge needle (Nanofil, World Precision Instruments) was lowered to the target depth, and the drug was infused at 0.1 µl/min. The needle was held in place for 3–5 minutes post-infusion before slow withdrawal (≤1 mm/min). Dental cement mixed with cyanoacrylate was used to cover infusion sites. Antibiotic cover (TMS, 1 mg/ml) was provided in the drinking water for 5 days post-surgery. Lesion extent was verified by brightfield imaging of 35 μm coronal sections counterstained with Neurotrace 500/525 fluorescent Nissl stain.

#### Chemogenetic Inhibition

##### CA1 PV Interneuron Inhibition

###### *Viral Infusion*

PV-T2A-Cre mice were infused with AAV5-hSyn-DIO-hM4Di (Gi)-mCherry (Addgene #44362; $1.4 \times 10^{12}$ vg/ml, 500 nl per hemisphere) bilaterally into dorsal CA1 (AP: −1.9 mm, ML: ±1.3 mm, DV: −1.3 mm from bregma) using a 32-gauge NanoFil syringe (World Precision Instruments) at 60 nl/min. The needle was left in place for 10 minutes post-infusion before withdrawal. Meloxicam (5 mg/kg, subcutaneous) was administered on the day of surgery and for five consecutive days thereafter (Figure 2e-g, Supplementary Figure 7, 8).

###### *Behavioral Protocol*

Three weeks after viral infusion, animals were handled and habituated for five days. Animals were then trained with the single-shock generalization paradigm in context A. Animals were tested remotely in ABC order, receiving an intraperitoneal injection of CNO (3 mg/kg; see CNO Preparation below) 45 minutes prior to context B testing on the CNO test day, and vehicle (7% DMSO in 0.9% saline, equivalent volume) prior to context A and context C testing. The same animals subsequently served as their own vehicle controls, receiving vehicle prior to context B in a second ABC sequence. Behavior was evaluated only from animals in which mCherry expression was confined to dorsal CA1, verified by post-hoc immunofluorescence (Figure 2e, f).

##### Retrosplenial Cortex Inhibition

###### *Viral Infusion*

C57BL/6J animals (n = 41, 2–3 months) were trained in context A using the three-shock generalization paradigm. Two days after training, a cocktail of AAV9-CaMKII-HI-GFP-Cre-WPRE-SV40 (Addgene #105551; $2.5 \times 10^9$ vg) and AAV5-hSyn-DIO-hM4Di(Gi)-mCherry (Addgene #44362; $6 \times 10^8$ vg) in a 1:1 ratio (500 nl per hemisphere) was infused bilaterally into

RSc (AP: −2.0 mm, ML: ±0.3 mm, DV: −0.7 mm from bregma). This combinatorial strategy restricts hM4Di expression to CaMKII-positive (excitatory) neurons of RSc. Meloxicam (5 mg/kg, subcutaneous) was administered for five days post-surgery (Figure 4a, b).

###### ***Behavioral Protocol***

These animals were then tested in ABC order beginning two weeks after viral infusion (Figure 4a). Context A was tested on day 30, context B on day 31, and context C on day 32. Vehicle (7% DMSO in 0.9% saline) was administered intraperitoneally 45 minutes prior to context A and context B testing. Animals were then randomly assigned by rank-order of context A freezing to ensure balanced group means, and received either CNO (3 mg/kg, IP) or vehicle 45 minutes prior to context C testing. After behavior completion, animals were transcardially perfused and brains were sectioned for verification of viral expression in RSc by fluorescence microscopy (Zeiss Observer.Z1, 10× and Zeiss LSM 880 Airyscan).

##### CNO Preparation and Administration

CNO powder (Sigma-Aldrich, catalogue #C0832-5MG) was dissolved in DMSO by gentle agitation until completely dissolved. Saline (0.9% NaCl) was added to achieve a final working concentration of 450 µg/ml (7% DMSO in saline). Working solution was prepared fresh for each experimental day and stored at room temperature. CNO was administered intraperitoneally at 3 mg/kg body weight, 45 minutes prior to the relevant behavioral session. Vehicle controls received equivalent volumes of 7% DMSO in 0.9% saline.

#### Histology and Immunohistochemistry

##### Perfusion and Tissue Preparation

Animals were anesthetized with ketamine (90 mg/kg) and xylazine (10 mg/kg) cocktail administered subcutaneously. Depth of anesthesia was confirmed by the absence of toe-pinch reflex. Animals were transcardially perfused with ice-cold 1× PBS followed by 4% paraformaldehyde (PFA) in PBS. Brains were post-fixed in 4% PFA overnight at 4°C, transferred to 30% sucrose in PBS, and stored at 4°C until fully equilibrated (brain sinks to bottom of tube). Brains were sectioned at 50 µm using a cryostat (Leica) and stored in 0.1% sodium azide in 1× PBS at 4°C until staining.

##### cFos Immunostaining (LaBCAM experiments)

Free-floating sections were incubated in blocking solution (5% goat serum, 0.3% Triton X-100 in 1× PBS) for 2 hours at room temperature. Sections were incubated with guinea pig monoclonal anti-cFos primary antibody (1:1700; Synaptic Systems, catalogue #226308) for 48 hours at 4°C in antibody dilution buffer (1% BSA, 0.3% Triton X-100 in 1× PBS). After three PBS washes (10 minutes each), sections were incubated with goat anti-guinea pig IgG secondary antibody conjugated to Alexa Fluor 647 (1:1000; ThermoFisher, catalogue #A-21450) for 2 hours at room temperature. Sections were washed, mounted on slides, and coverslipped with DAPI-Prolong antifade fluoromount solution. Sections were imaged on a Zeiss Axio Scope at 10× (NA 0.4) and 20× magnification (Figure 3a).

##### Parvalbumin Immunostaining (PV-IN experiments)

Free-floating sections were blocked (5% goat serum, 0.3% Triton X-100 in 1× PBS, 2 hours) then incubated overnight with rabbit polyclonal anti-PV primary antibody (1:1500; ABclonal, catalogue #A2791) in antibody dilution buffer (1% BSA, 0.3% Triton X-100 in 1× PBS). After three PBS washes (10 minutes each), sections were incubated with goat anti-rabbit IgG $Fab_2$-Alexa Fluor 488 secondary antibody (1:1000; Cell Signaling Technology, catalogue #4412S) for 2 hours at room temperature. Sections were washed, mounted, and coverslipped with Vectashield antifade mounting medium with DAPI (Vector Laboratories, H-1200). Images were acquired on a Zeiss Observer.Z1 inverted fluorescence microscope at 10× (NA 0.45, Plan-Apochromat M27).

#### Brainwide Correlated Activity Mapping (LaBCAM)

##### Tissue Collection and Staining

C57BL/6J mice (2–3 months) were trained in the three-shock generalization paradigm and tested remotely in either ABC or BAC order. Animals were transcardially perfused 90 minutes after context C testing on day 33. Brain sections (50 μm) were processed for cFos immunostaining as described above.

##### Region of Interest Identification and Cell Counting

Stained sections were imaged and anatomically registered to the Allen Mouse Brain Atlas using the BigWarp plugin in ImageJ/Fiji to isolate regions of interest. Twenty brain regions were analyzed, spanning cortical, hippocampal, thalamic, and limbic areas. A random forest classifier was trained using pixel and object classification in ILASTIK (Interactive Learning and Segmentation Toolkit) to identify cFos-positive cells. The output CSV file contains object class (positive, negative, merged, background), count, intensity, classification probability, and XYZ coordinates. Post-processing filters based on object size and classification probability were applied to retain only true cFos-positive cells (Figure 3, Supplementary Figure 10, 11).

**Correlated Activity Analysis and Network Visualization**

Median cFos fluorescence intensity was calculated per region per animal. Pairwise Pearson correlation coefficients were computed across animals for all region pairs using OriginPro software (ver. 10). Correlation matrices were constructed separately for ABC and BAC groups. Network graphs were visualized using RStudio, with nodes representing brain regions (size proportional to median cFos intensity) and edges representing pairwise correlations (edge thickness and transparency proportional to $|r|$ value; red = positive correlation, blue = negative correlation). Only correlations meeting the threshold $r \geq +0.5$ or $r \leq -0.5$ were displayed. Brain regions were color-grouped by major subdivision: mPFC (ACC, PL, IL), RSc (RSA, RSG), dorsal hippocampus (dCA1, dCA2, dCA3, dDG), ventral hippocampus (vCA1, vCA2, vCA3, vDG), amygdala (CeA, BLA), anterior thalamus (ACM, ANRE), posterior thalamus (CM, NRE, PVT).

*Note: Correlated cFos intensity across animals reflects co-activation patterns and is used here as a proxy for functional co-engagement of brain regions during HOA retrieval. This approach does not establish direct anatomical connectivity between regions.*

#### *In Vivo* Two-Photon Imaging of Dendritic Spines

**Animals and Craniotomy**

Thy1-EYFP mice (8–10 weeks) underwent craniotomy for implantation of a chronic cranial window over RSc. Under isoflurane anesthesia (2%), carprofen (150 µl, 1 mg/ml, subcutaneous) and dexamethasone were administered. After hair removal and scalp incision, a 5 mm diameter craniotomy was made over RSc using a pneumatic drill and a circular cover glass was fixed over intact dura using cyanoacrylate adhesive and dental cement. An aluminum head bar was fixed rostral to the imaging window. Animals were allowed to recover for approximately two weeks before training.

**Imaging Protocol**

Baseline two-photon imaging was performed 24–48 hours before remote testing in ABC or BAC order (approximately 28 days post-training). Longitudinal imaging sessions were performed after ABC or BAC remote testing (Figure 4e, Supplementary Figure 12). For each session, animals were anesthetized with isoflurane (2-3% with room air @ 150-200ml/min during induction; 0.5-1% @ 70-100ml/min during maintenance for *in vivo* imaging)

Animals were head-fixed on the imaging stage and body temperature was maintained at 37°C using a warm pad. Cranial windows were cleaned with sterile water before each session. Brightfield images of the cortical vasculature were acquired using blue LED illumination before each session to align fields of view across sessions. Two-photon imaging was performed on a custom-modified Zeiss Axio Examiner Z.1 using an ultrafast pulsed Ti:Sapphire laser (Tsunami, Spectra Physics) tuned to 910 nm. Power at the back aperture of the objective was set to 150–200 mW (Olympus XLPLN25XWMP2, 25×, NA 1.05, water-immersion). Images were acquired by raster scanning using a 2D galvanometer mirror system (GVSM002, Thorlabs). Image acquisition was controlled by ScanImage 3.8 software integrated with a data acquisition card (NI-PCI 6110). Z-stacks were acquired at 1 μm steps (100–150 frames per stack) at 1024 × 1024-pixel resolution (4 ms/line). Signal was collected via a low-noise current pre-amplifier (SRS-570) in high-bandwidth mode (800 kHz, sensitivity 10 μA/V). Image stacks were saved as 16-bit TIFF files (Figure 4e, Supplementary Figure 13).

**Dendritic spine counts**

Dendritic spines were traced and quantified using the Simple Neurite Tracer (SNT) plugin in Fiji. Dendrites were manually traced and each spine was marked as a branch point. Dendrite length in three dimensions was estimated automatically by the software; the number of branch points reflects spine count per dendrite. Spine coordinates were exported for raw spine counts and preliminary clustering estimates. Since raw counts with such tracing do not capture 3-D geodesic aspect of the spine dynamics, we then performed spine segmentation and clustering analyses using machine learning-based approach.

#### Spine Segmentation and Clustering Analysis

We trained and applied a random forest classifier-based machine learning approach (using the Interactive Learning and Segmentation Toolkit, Ilastik) to automatically detect dendrites and spines in raw images and quantify their changes across five imaging sessions. The ML pipeline began with pixel classification, training the model to assign each pixel to one of four categories: dendrite, spine, background, or out-of-focus. Prediction maps from this step were then used to generate two refined subclasses: "dendrite with spine" and "spine only." These maps were further used for object classification, during which the system was trained to segment structures into "spine" and "dendrite with spine" classes. This yielded unique identification numbers (IDs) for individual dendrites, along with the spines attached to each, including their geodesic distances and XYZ coordinates. From the geodesic distances, we computed nearest-neighbor distance (NN) and inter-spine distance (ISD; i.e., number of spines per unit dendrite length). Analyses were restricted to dendrites with at least three spines. (Supplementary Figure 14, Figure 4f-h).

We quantified the average inter-spine distance $\langle \delta_{ISD} \rangle$ and average nearest-neighbour distance $\langle \delta_{NN} \rangle$ for each dendrite following object classification. As a metric of spine clustering, we computed the extent (ε) of clustering as the difference $\langle \delta_{ISD} \rangle$ - $\langle \delta_{NN} \rangle$ and divided it by the average inter-spine distance $\langle \delta_{ISD} \rangle$ per dendrite. i.e.

$$\varepsilon_{clustering} = \left( \frac{\langle \delta_{ISD} \rangle - \delta_{NN}}{\langle \delta_{ISD} \rangle} \right)$$

We note, since we are measuring the nearest neighbor distance, that this quantity ($\varepsilon_{clustering}$) would be inherently < 1 on an average. In order to capture the relative change in clustering dynamics we normalize this quantity ($\varepsilon_{clustering}$) with its baseline value for respective group (ABC and BAC), as estimated for imaging session conducted before the remote-retrieval tests (ABC or BAC), and refer to it as relative extent of spine clustering (Relative $\varepsilon_{clustering}$). Thus, a reduction in this quantity implies an increase in nearest neighbor distance ($\delta_{NN}$) and a reduction in the extent of clustering (Relative $\varepsilon_{clustering}$).

Comparing spine clustering between the ABC (HOA) and BAC (control) groups revealed that clustering reverted to baseline levels in the BAC group when imaged few days after CtxC testing, whereas it persisted in the ABC group up to ~3-weeks post-testing. We attribute this difference to the role of the Retrosplenial cortex (RSc) in conflict resolution, and that being consolidated specifically for the ABC order.

#### APP/PS1 and Aged Animal Experiments

##### APP/PS1 Alzheimer's Disease Model

APP/PS1 double-transgenic mice (JAX #034829; expressing Mo/HuAPP695swe and PS1-dE9 under neuronal promoters) and wild-type littermates of the same C3H genetic background were used. Both male and female animals aged 2–5 months were used, prior to the onset of amyloid plaque deposition (which begins at 6–7 months in this line). Animals were trained in the three-shock generalization paradigm and tested remotely in either ABC or BAC order on days 31, 32, and 33. Testing sessions on days 31 and 32 were conducted on the same day separated by 7 hours; context C testing on day 33 was followed by perfusion 90 minutes later for cFos analysis.

##### Aged Animals

Male C57BL/6J mice aged 16–17 months were trained in the three-shock generalization paradigm and tested remotely in ABC (n = 12) or BAC (n = 16) order on days 31, 32, and 33, using the same protocol as young adult animals.

#### Behavior Analysis

All behavioral sessions were video-recorded via a back-mounted camera at 25 fps and converted to AVI format using Avidemux 2.6 (64-bit; MPEG video compression, constant frame rate). Freezing behavior was defined as the absence of all movement except breathing. Converted videos were opened in ImageJ using a custom semi-automated plugin (Scoring_assistant). Scoring was performed manually at a resolution of 10 frames per click (0.4 seconds per scored window). For each window, a left click recorded freezing (1 = no movement except breathing) and a right click recorded movement (0). Scoring began from the frame at which the chamber door closed after animal release and ended when the door opened for animal removal. The binary score column was processed using a custom Excel macro that filtered for bouts of three or more consecutive freezing responses, corresponding to a minimum continuous freezing duration of 30 frames (1.2 seconds).

Percentage freezing was calculated from the filtered scores over 120 seconds of context exposure. Percentage time freezing was calculated for each testing session and used as the primary behavioral measure. Animals with viral expression outside target regions, failed lesions, or atypical freezing profiles (<10% freezing in training context A at remote testing) were excluded from analysis.

#### Supplementary Figures

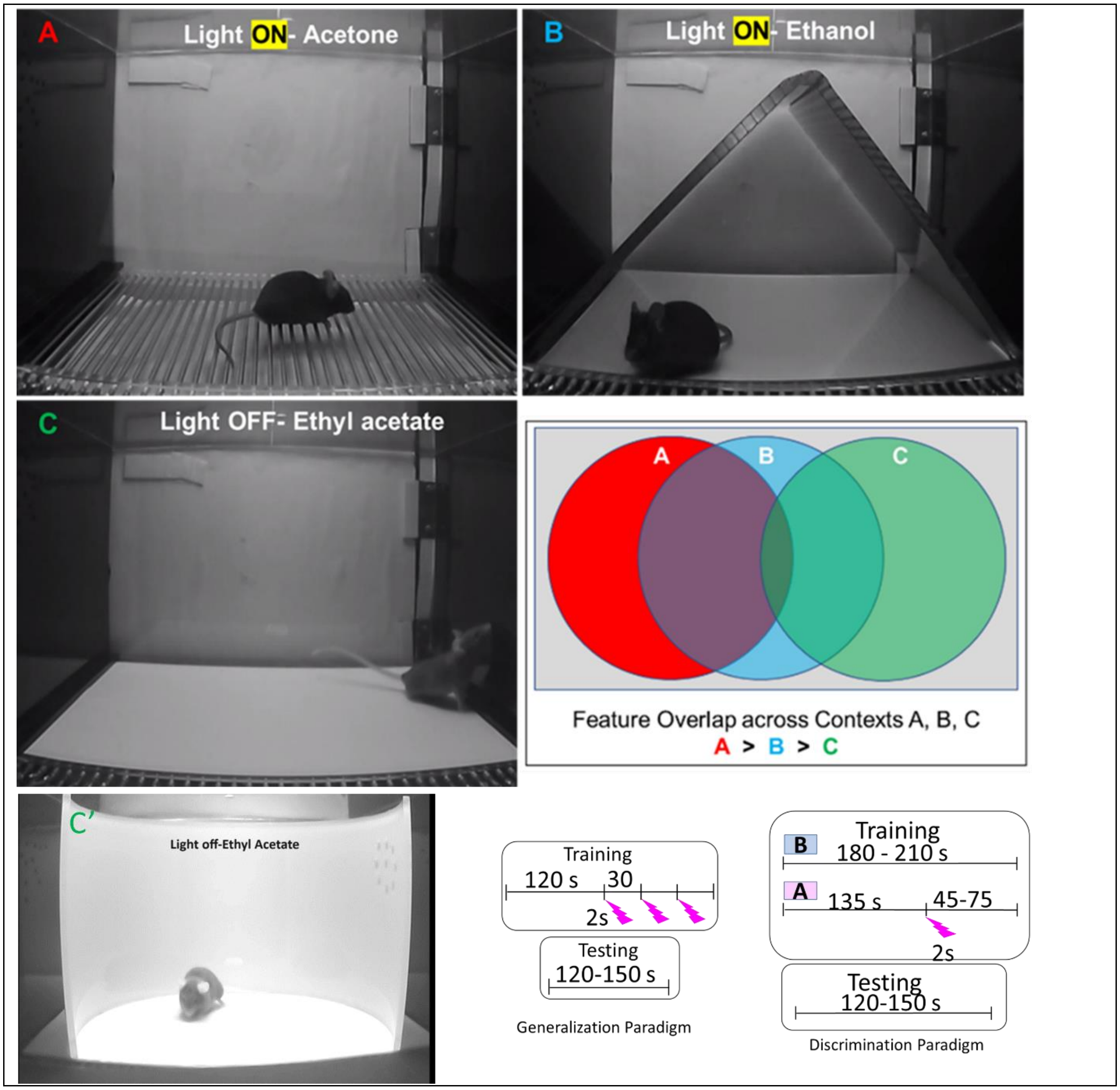


**Supplementary Figure 1. Four contexts (A, B and C, C') used in the study.** Venn diagram to depict the decreasing feature overlap across the contexts A, B and C. Contexts C and C' showed similar results and hence are used interchangeably in the manuscript. The training protocol for generalization paradigm shows 120s long context exposure followed by three shocks at 30 s intervals with last 30s as post shock exploration. Training protocol for discrimination paradigm shows morning exposure to context B for 180-210 s, 6-hours later, same animals are exposed to context A with 135 s exploration before one shock followed by 45-75s post-shock exploration. This training is repeated across three days. Testing for bpth generalization and discrimination paradigms was similar with free exposure to contexts for 120-150s at defined testing intervals.

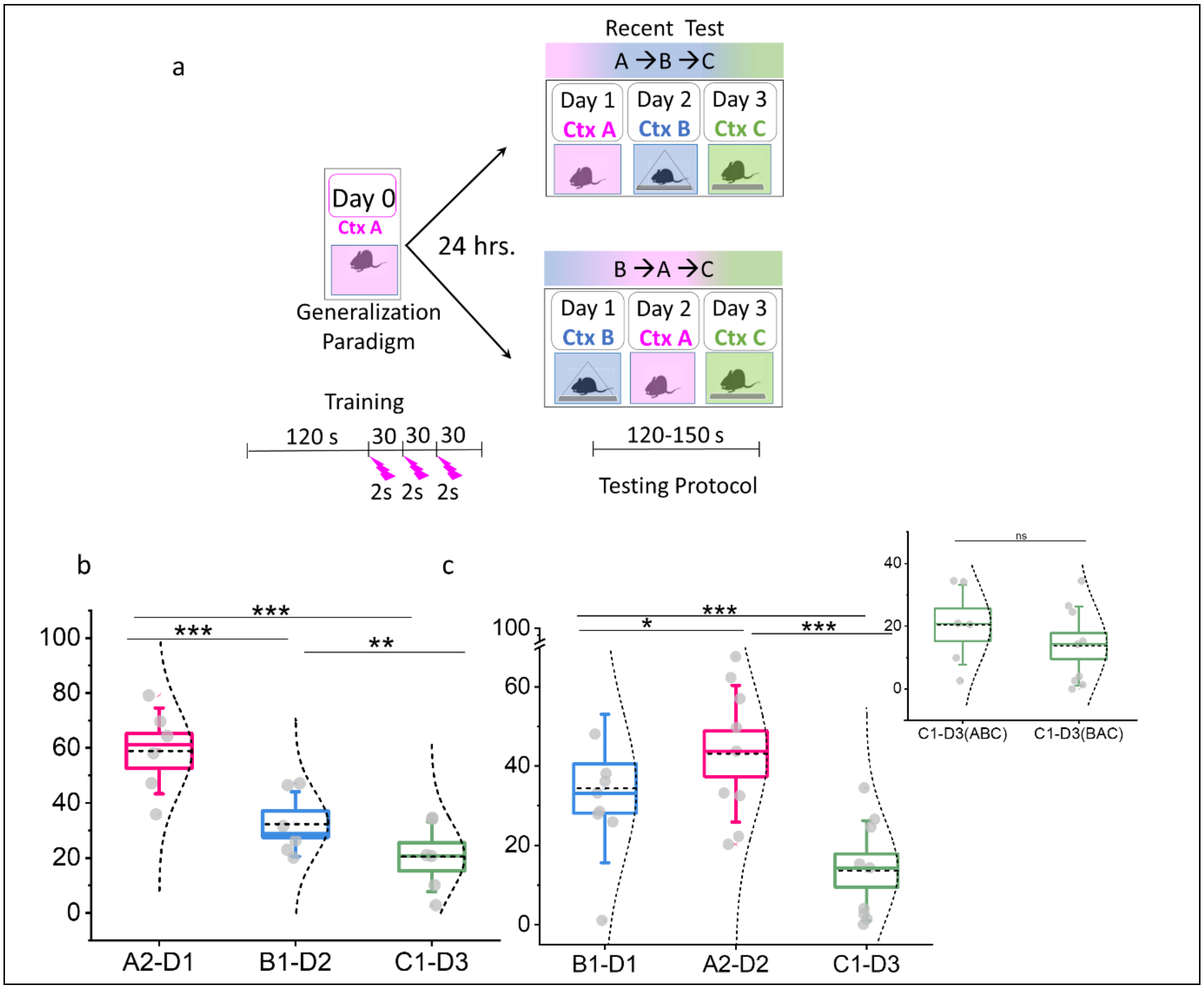


**Supplementary Figure 2. Formation of HOA is exclusive to remote memory.** a) Animals were trained in three shocks and tested either in ABC or BAC order. Animals showed contextual discrimination when tested in (b) ABC order (1w-RM ANOVA, $F_{2,4}$=176.98, P>F=1.25E-4), and (c) BAC order (1w-RM AVONA, $F_{2,7}$=53.94, P>F=5.58). Animals from both groups showed higher freezing in the training context, CtxA, than CtxB and CtxC. However, (inset) no significant difference in context C freezing was observed when comparing C1-D3(BAC) and C1-D3(ABC) one-tailed Two sample t-test ($|t|_{13}$=1.01, P>|t|=0.32), suggesting that formation of HOA is a remote exclusive phenomenon.

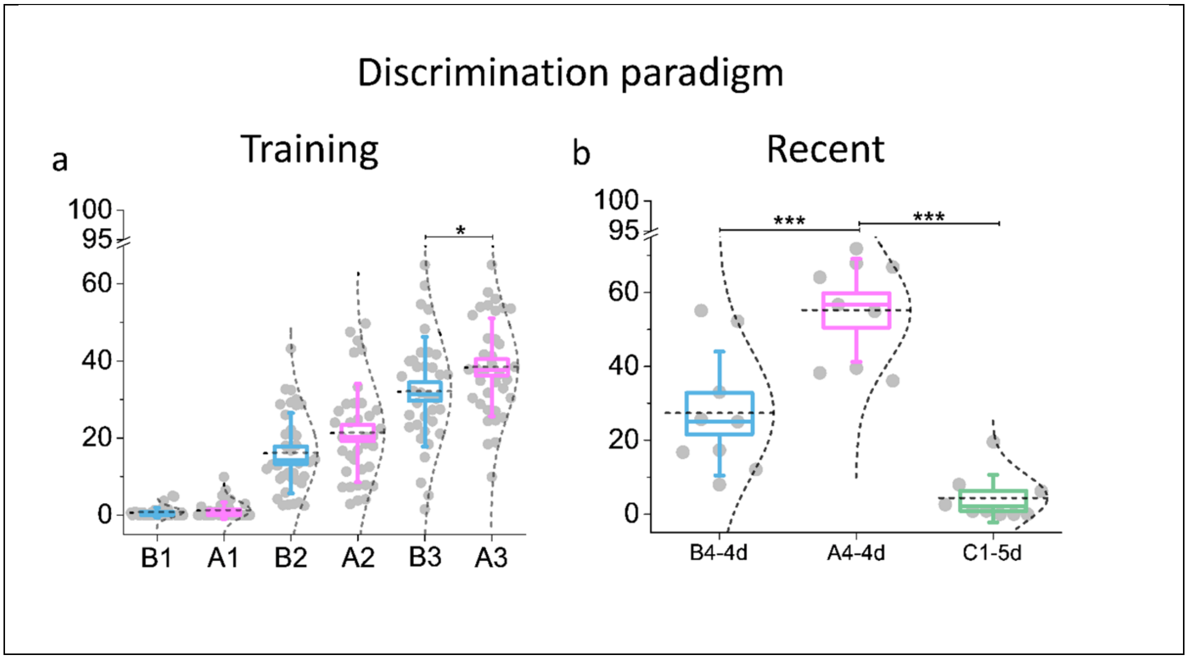


**Supplementary Figure 3. Early freezing response in discrimination paradigm.** a) Progressively increasing freezing response in context B and A during training, leading to significantly higher fear recall in A than B on day 3 (1-w RM ANOVA, $F_{5,31} = 67.15$, P>F = 1.06E-15, n = 36; Post-hoc Tukey test B3 vs. A3: $|t|_{42} = 4.39$, P > |t|= 0.0267). b) Recent test for discrimination memory shows significantly higher freezing in context A vs. B on day 4, with significantly lower freezing in C on day 5.

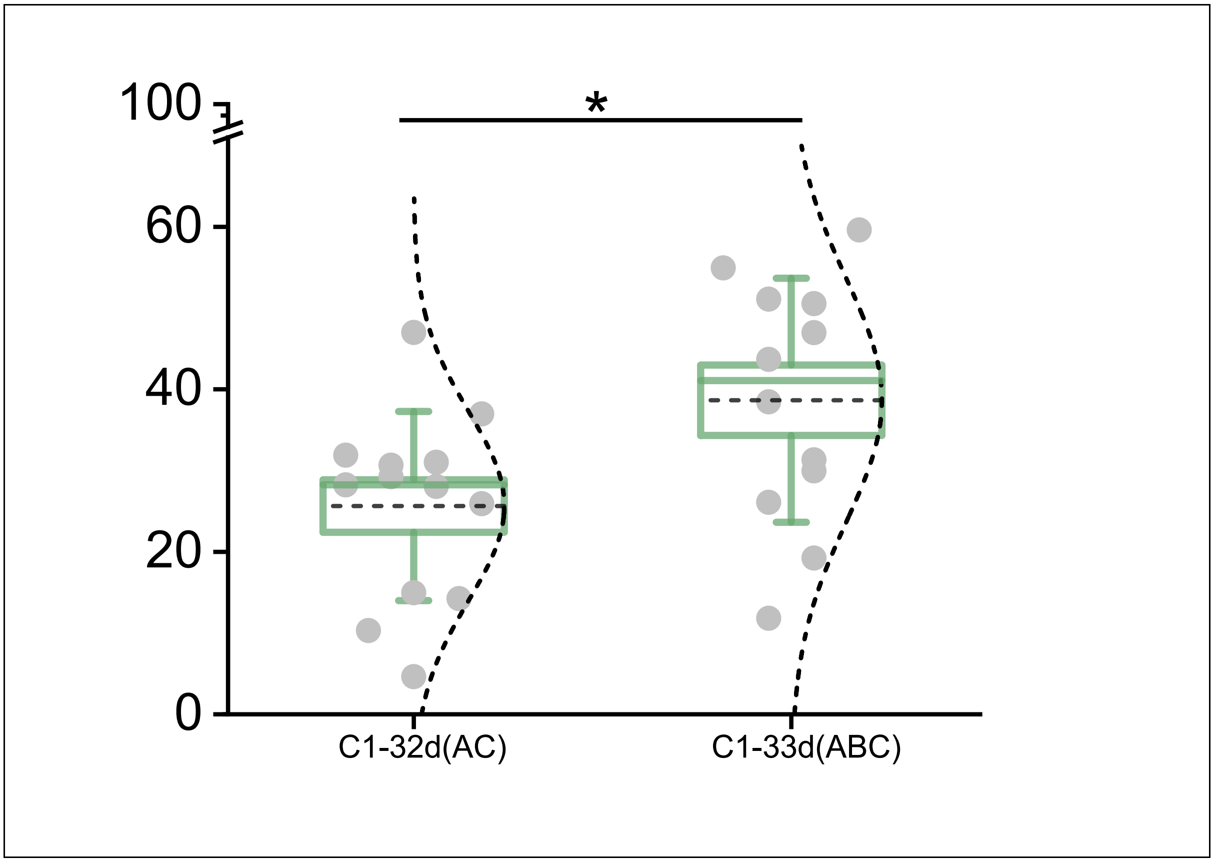


**Supplementary Figure 4.** Context C freezing in absence of intervening exposure to context B (C1-32d, Figure 1f) is significantly lower than the context C freezing during ABC testing leading to HOA based fear expression (C1-33d, Figure 1c)

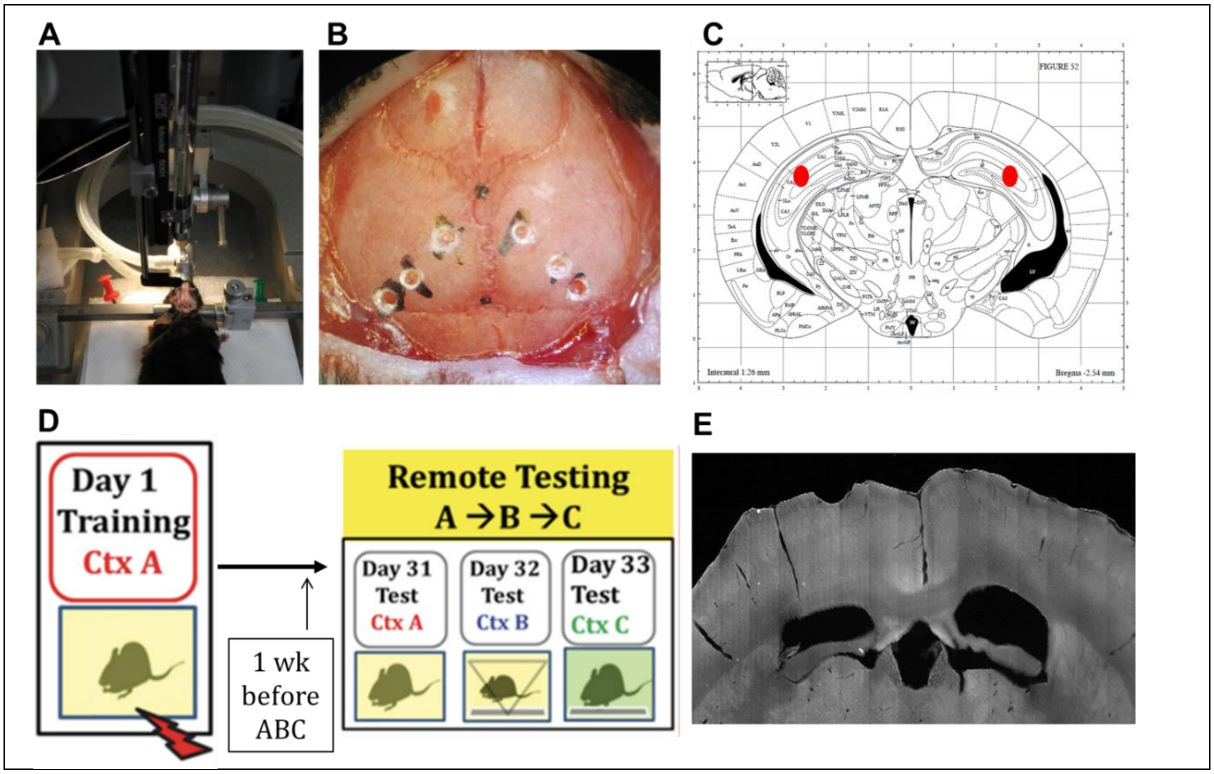


**Supplementary Figure 5.** Hippocampal lesion with ibotenic acid infusion.

**a)** Anesthetized mouse mounted on stereotax for drug infusion.

**b)** Mouse skull marked with black dots for bregma (midline top) and lambda (midline bottom). Infusion sites are marked as explained above and drilled-open with pneumatic drill. The opening is made with slightly larger diameter than the needle. Infusion sites: Infusion Site Pair 1 = Rostro-caudal from Bregma: - 1.3, Mediolaterally: ± 1.2; Infusion Site Pair 2 = Rostro-caudal: - 2.5, Mediolaterally: ± 2.5; Infusion Site Pair 3 = Rostro-caudal: - 3.3, Mediolaterally: ± 3.1.

**c)** Mouse coronal diagram showing one of the representative drug infusion sites for hippocampus.

**d)** Timeline for hippocampus lesion with respect to training and remote testing. Lesions are performed three weeks after training and animals are allowed to recover for 1 week before testing them in ABC order. Two control sub-groups are infused with drug vehicle (1x PBS, pH 7.4) at the same time and allowed to recover for 1 week. One control sub-group is tested in ABC testing order, while the other control sub-group is tested in BAC testing order.

**e)** One of the coronal brain sections showing dorsal hippocampal lesion from experimental mice.

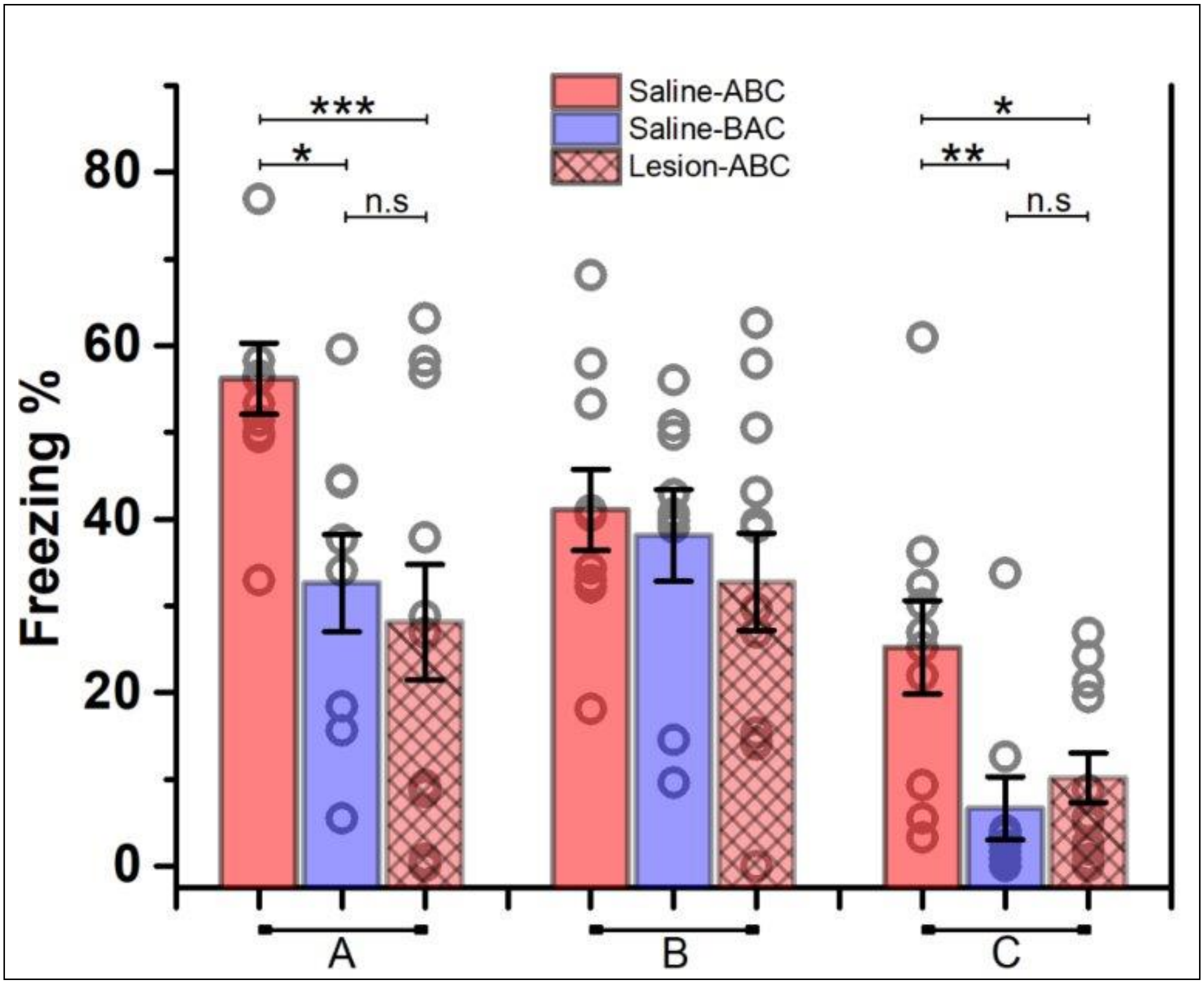


**Supplementary Figure 6.** Comparison of freezing across three groups of animals injected after three weeks of training in context A: Saline-ABC group was injected with saline and tested in ABC order, Saline-BAC group was injected with saline and tested in BAC order, and Lesion-ABC group was injected with ibotenate and tested in ABC order. Comparing the freezing in CtxA across the three conditions, freezing for A-Sal-ABC (56.20 ±4.11) was higher than A-Sal-BAC (32. 63±5.60), and A-Lsn-ABC (28.11±6.63). The difference was significant between control (saline) animals tested in A vs. lesioned animals tested in ABC or saline animals in BAC ($F(2,28) = 6.92$, $P>F = 0.004$; Bonferroni's Post hoc: A-sal-BAC vs. A-sal-ABC – $t = -2.77$, $p>t = 0.03$; A-Lsn-ABC vs. A-Sal-ABC – $t = -3.54$, $p>t = 0.004$; A-Lsn-ABC vs. A-Sal-BAC – $t = -0.55$, $p>t$ = n.s). Freezing for context B was found similar across the three conditions (B-Sal-ABC: 41.06±4.67, B-Sal-BAC: 38.12±5.29), B-Lsn-ABC: 32.76±5.61; $F(2,28) = 0.67$, $P>F$ = n.s). However, freezing in context C showed HOA-based increase in freezing only for saline injected animals tested in ABC with significantly lower relative freezing in the other two conditions. (C-Sal-ABC: 25.21±5.39, C-Sal-BAC: 6.65±3.62, C-Lsn-ABC: 10.16±2.85; $F(2,28) = 5.81$, $P>F = 0.008$; Bonferroni's Post hoc: C-sal-BAC vs. C-sal-ABC – $t = -3.14$, $p>t = 0.012$; C-Lsn-ABC vs. C-SalABC – $t = -2.73$, $p>t = 0.032$; C-Lsn-ABC vs. C-Sal-BAC – $t = 0.62$, $p>t$ = n.s). Tukey's post hoc also gave similar results.

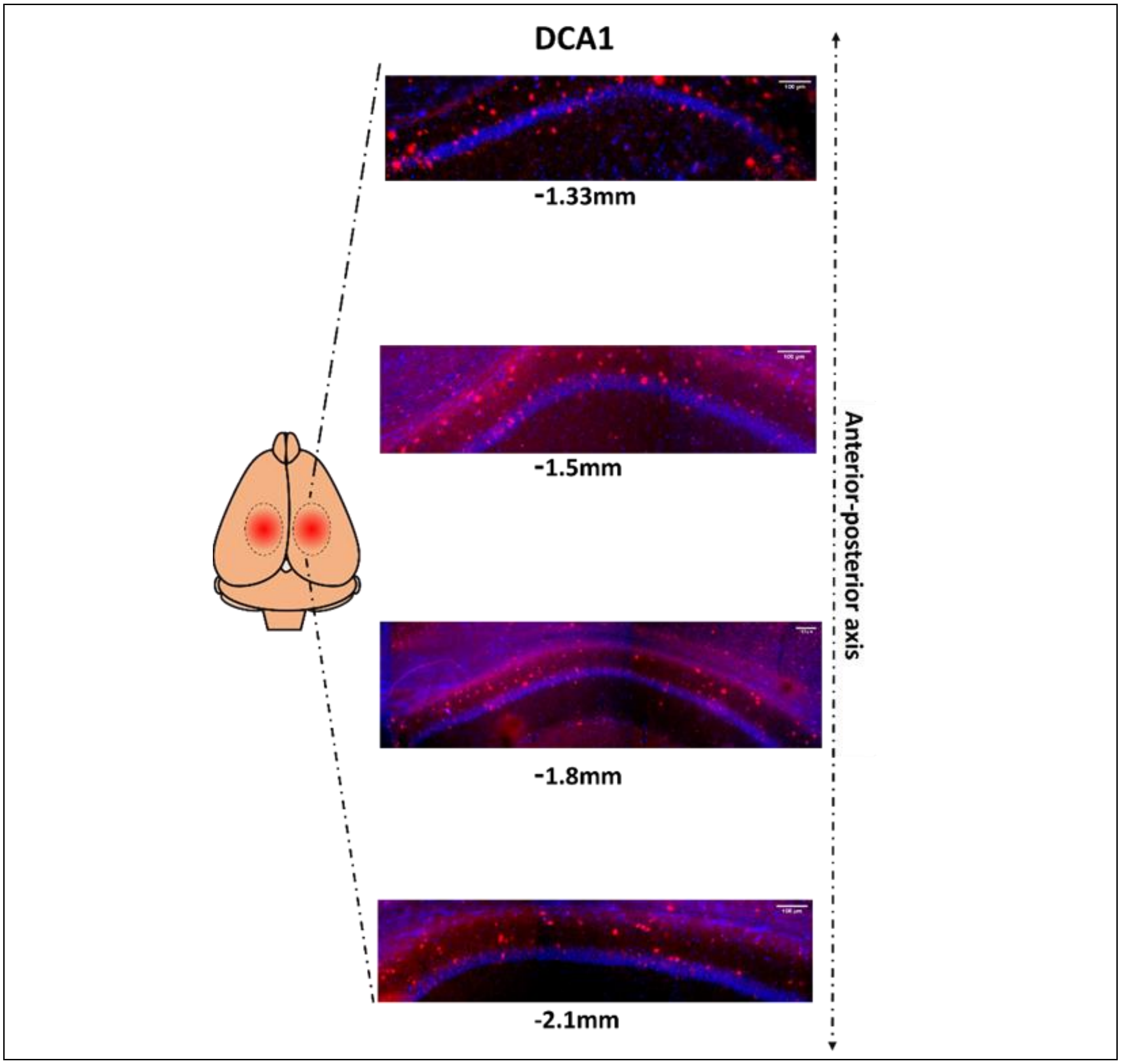


**Supplementary Figure 7. Virus expression in the dHpC along the rostro-caudal axis**
After completion of the behavioral experiments, animals were transcardially perfused, and brains were collected. Tissues were then cryopreserved and sectioned at thickness of 50um using microtome, followed by imaging the sections using inverted fluorescence microscope (Zeiss, Observer.Z1) at 10X (NA:0.45, Plan Apochromat M27) magnification. The expression of the AAV5-hSyn-DIO-hM4Di (Gi)-mCherry virus was checked along the Rostro-Caudal axis (starting from –1.33mm from bregma to -.2.1 mm).

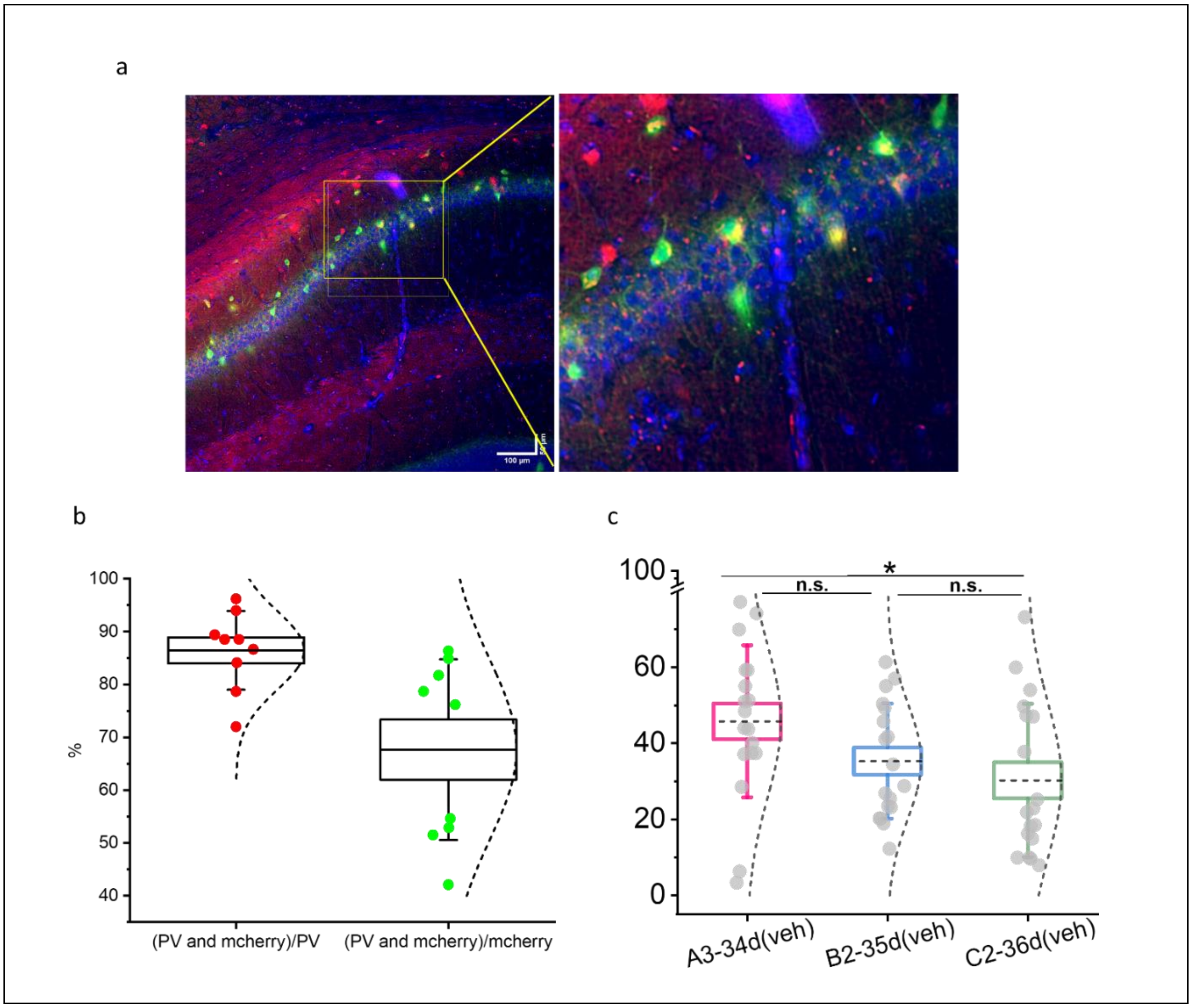


**Supplementary Figure 8.** DREADD expression and effect of chemogenetic inhibition of PV interneurons on conditional suppression of HOA acquisition.

a) DREADD expression after AAV infusion in dorsal hippocampus and overlap with parvalbumin (PV) interneurons. Green – PV antibody immunolabelled neurons, Red – mCherry expression in AAV infected neurons, Yellow – colocalization of green and red neurons (PV expressing AAV infected neurons).

b) After completion of the behavioral experiment, brains(n=5) were extracted and sectioned 50 µm thick and looked for the expression of inhibitory DREADD and its colocalization with PV-IN. For analysis, 1-2 sections from each animal were taken, ranging from -1.2 mm to -2.1 mm (posterior to bregma). 86% endogenous PV-IN expressed hM4D(i)-mCherry, translating to high transfection efficiency, where 68% of mCherry expressed in PV-IN, indicating high selectivity.

c) Veh treated group (PV inhibition): Animals were further retested in ABC order, but in the presence of vehicle(7% DMSO in saline), and animals showed the highest freezing in A3-34d(VEH)(45.72±4.71) than that of C2-36d(VEH)(30.20±4.75) (1w-RM ANOVA, $F_{2,16}$=4.08, P>F=0.037) and posthoc Tukey's test ($|t|_{34}$=4.63, P>|t|=0.007). Where B2-35d (35.28±3.56) is not significantly different from C2-36d (posthoc Tukey's test, $|t|_{34}$=1.51, P>|t|=0.54).

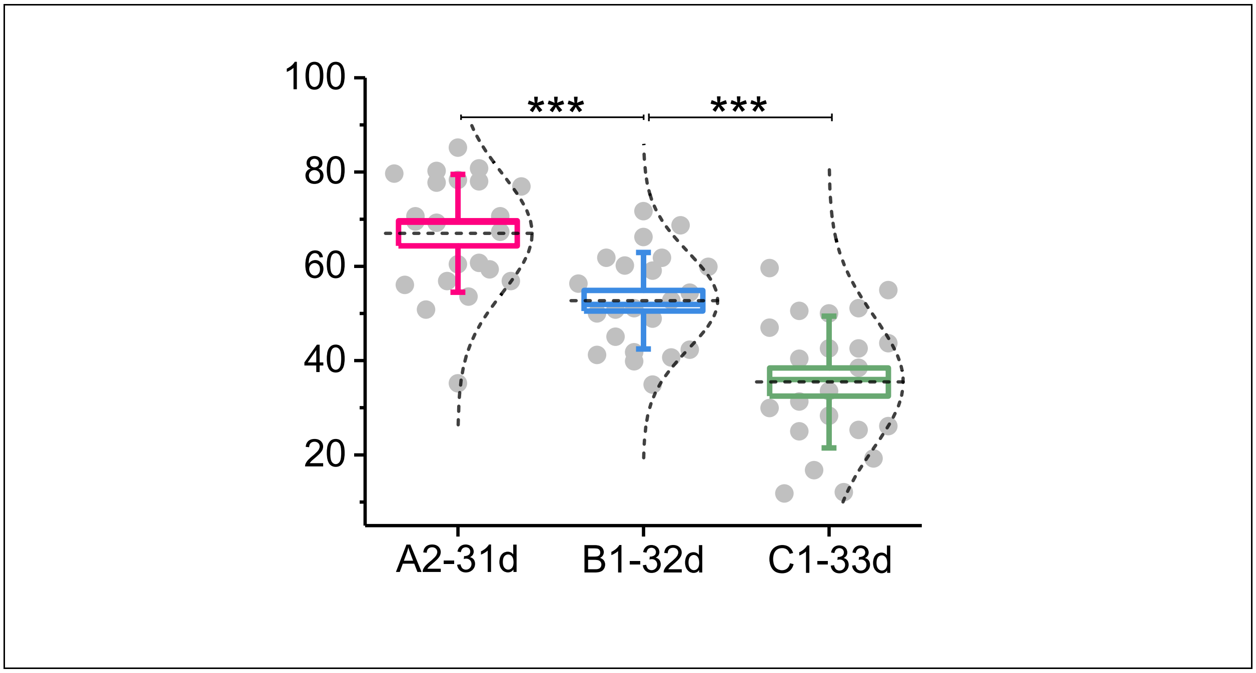


**Supplementary Figure 9.** Freezing during Remote ABC test before hippocampus lesions. Subset of these animals were tested for C-L-41d test.

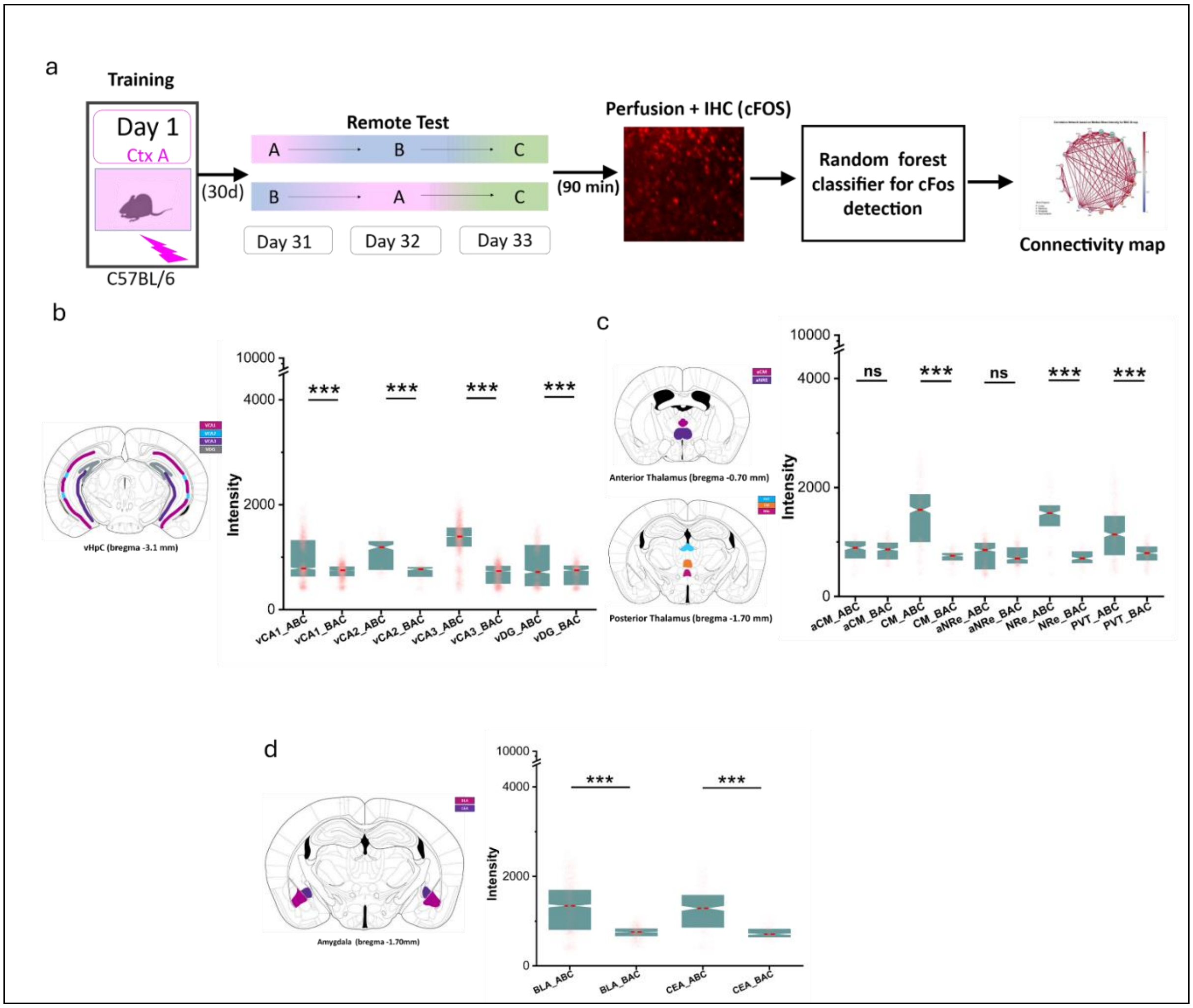


**Supplementary Figure 10.** LaBCAM – additional brain regions

a) Schematic Overview of Experimental Procedure: After contextual fear conditioning (CtxC), animals were perfused, and brain sections from the Ventral Hippocampus, Thalamus, and Amygdala were immuno-stained for cFos. A random forest classifier was employed to detect cFos-positive cells, enabling whole-brain activity mapping and the construction of a connectivity map across these regions.

**b) Ventral Hippocampus Subregions**: Subregions CA1, CA2, CA3, and DG exhibited significantly elevated mean cFos intensity when animals were tested in ABC order, as determined by the Kruskal-Wallis test with Dunn's post hoc analysis (VCA1: mean rank difference = 813.85, Z = 16.31, P = 8.24E-60; VCA2: mean rank difference = 169.53, Z = 11.94, P = 6.69E-33; VCA3: mean rank difference = 3460.14, Z = 64.49, P = not provided; VDG: mean rank difference = 268.34, Z = 8.10, P = 5.13E-16).

**c) Thalamus Subregions**: The anterior Centro-median nucleus (aCM) and anterior nucleus reuniens (aNRE) showed comparable cFos expression levels (aCM: mean rank difference = 3.20, Z = 0.22, P = 0.822; aNRE: mean rank difference = 14.00, Z = 0.677, P = 0.49). In contrast, other subregions, including the Centro-median nucleus (CM), nucleus reuniens (NRe), and paraventricular nucleus (PVT), displayed significantly higher cFos expression (CM: mean rank difference = 322.54, Z = 18.72, P = 3.02E-78; NRe: mean rank difference = 153.02, Z = 13.46, P = 2.54E-41; PVT: mean rank difference = 270.81, Z = 14.08, P = 4.71E-45).

**d) Amygdala Subregions**: Both the basolateral amygdala (BLA) and central amygdala (CEA) exhibited notably higher cFos expression (BLA: mean rank difference = 825.45, Z = 26.86, P = 5.04E-159; CEA: mean rank difference = 247.29, Z = 15.60, P = 7.20E-55).

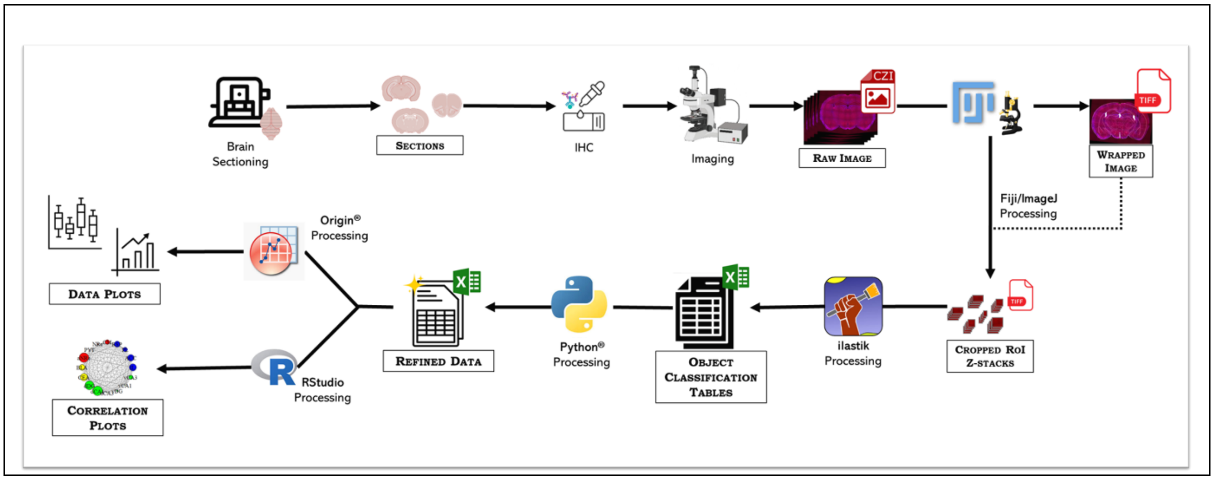


**Supplementary Figure 11.** Flowchart illustrating cFos detection and connectivity map creation: Brain sections from various regions were immuno-stained and imaged for cFos immediate early gene (IEG) expression. These sections were then aligned with corresponding moving images from a mouse brain atlas to identify and extract specific regions of interest (ROIs). The cropped images were batch-processed using a pre-trained random forest classifier, Ilastik, to detect cFos-positive cells. The output .csv file from the classifier included data on predicted classes (positive cell, negative cell, no object, merged positive cell, merged negative cell), along with their probabilities, size, intensity, and XYZ coordinates. This data was filtered based on size and probability to isolate true cFos-positive cells, which were then used to quantify cFos expression (intensity) in specific brain regions and to construct a connectivity map.

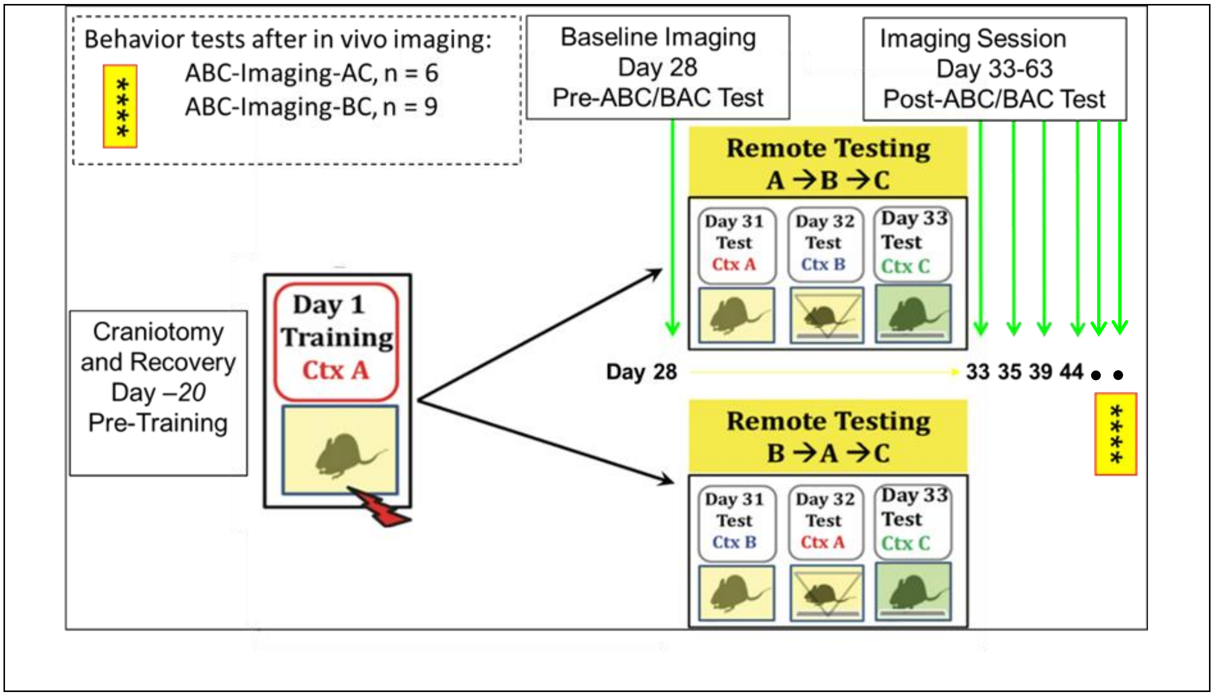


**Supplementary Figure 12.** Timeline showing training and testing with respect to craniotomy and *in vivo* imaging sessions. Craniotomy is performed ~20 days before training and mice are allowed to recover completely before training in context A. Baseline imaging is performed ~4 weeks after training. Experimental animals are tested in ABC order, control group is tested in BAC order. Longitudinal *in vivo* imaging is performed for six sessions after remote tests (green downwards arrows). Day for each session is mentioned below respective green arrow w.r.t. training day.

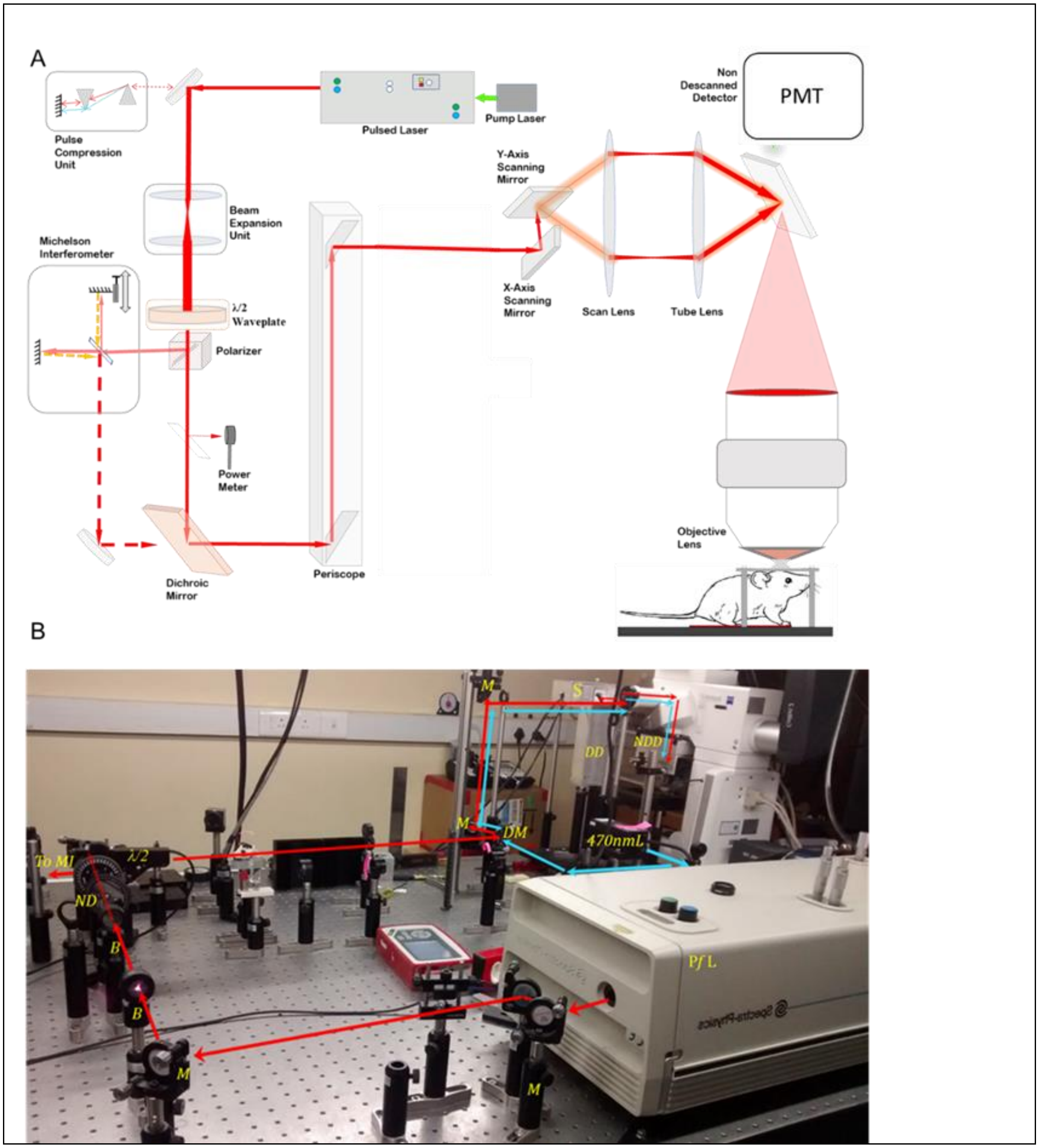


**Supplementary Figure 13.** Figure M8. Custom-built two-photon microscope. Top: Representative optical layout of customized two photon microscope set up for the *in vivo* imaging. (Not to scale) Bottom: Imaging set up in our lab. PfL-Pulsed femtosecond-laser, M-Mirror, B-Beam expansion unit, ND-Neutral density filter, λ/2-Half waveplate, DM-Dichroic Mirror, DD-Descanned Detector, NDD-Non-descanned detector, S-Scanner, 470nmL-470nmLaser, To MI – To Michelson Interferometer. Red Line- Optical path of pulsed laser for two photon excitation, Blue line-optical path of continuous wave laser for one photon excitation.

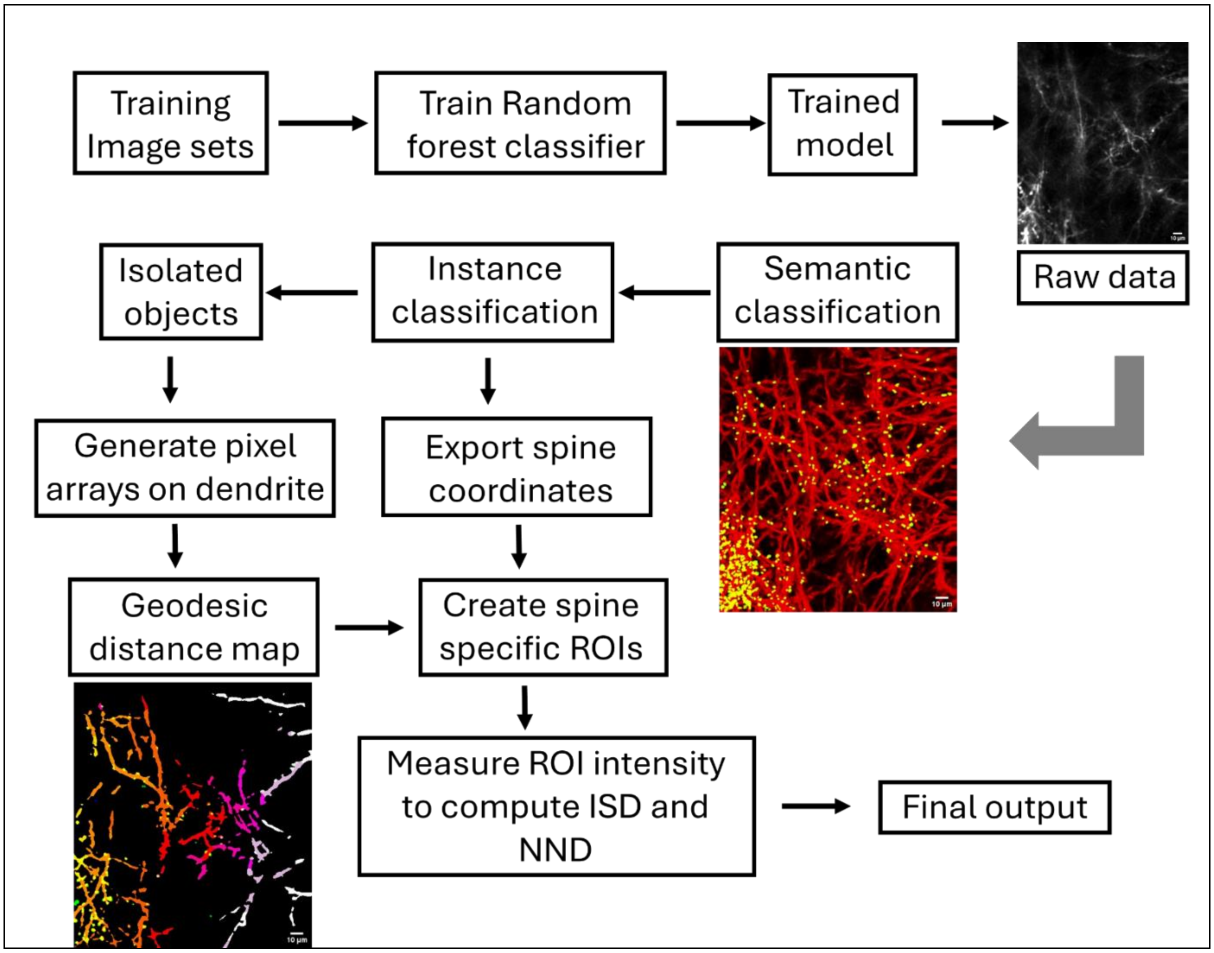


**Supplementary Figure 14**. Schematic illustration of spine reorganization analysis from RSc: A random forest classifier-based machine learning model (ilastik) was trained to segment dendrites and spines from raw 2-photon images. The system first assigns pixels to two broad classes, dendrite and spine, via semantic segmentation, followed by instance segmentation to isolate individual dendrites and spines as unique objects with their spatial coordinates. These coordinates were then used to generate a geodesic map, enabling the calculation of inter-spine distance (ISD) and nearest neighbor distance (NND) to assess dendritic spine reorganization.

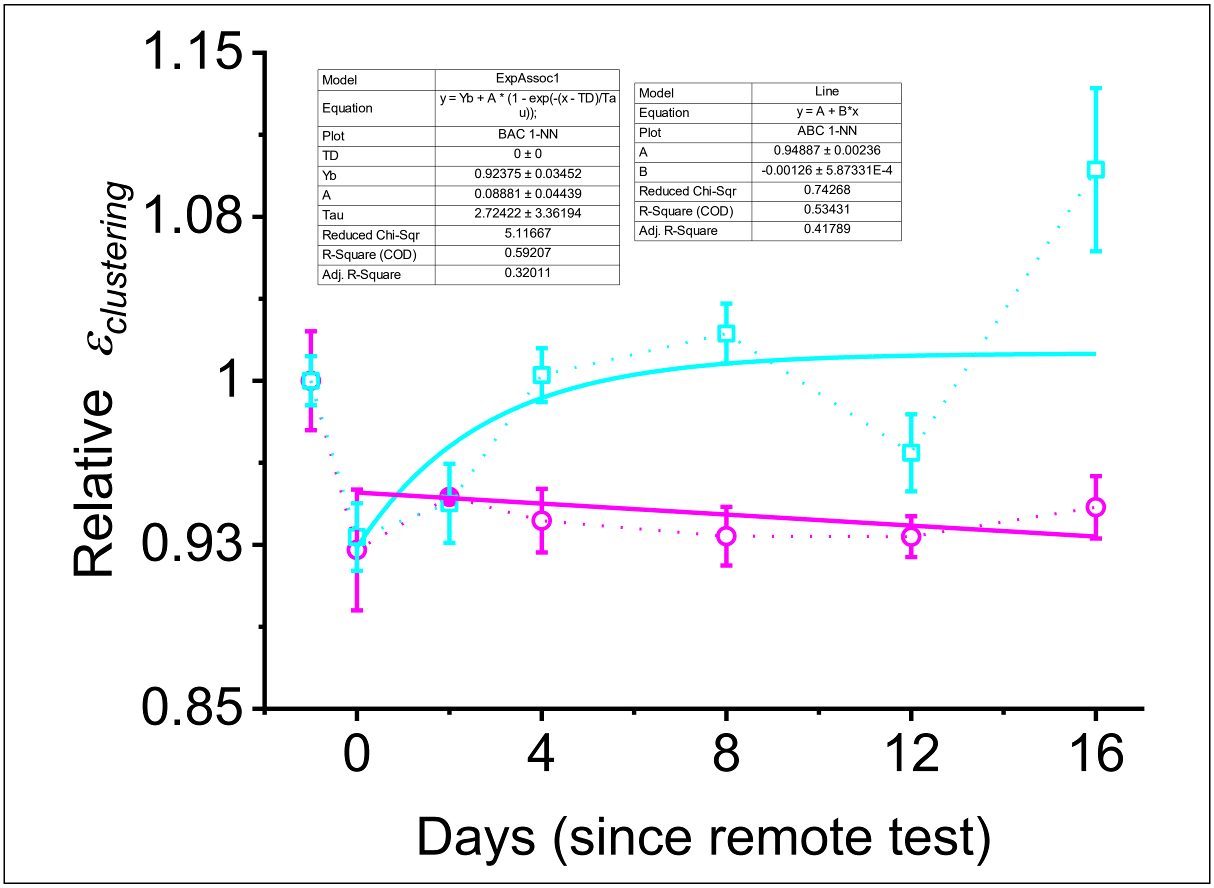


**Supplementary Figure 15. Dendritic spine reorganization following remote-recall mediated HOA-formation** (Circles, Magenta, ABC order) vs (Squares, Cyan, BAC order). Each datapoint indicates the Relative extent of clustering, days after HOA acquisition test (C-test - day 0). Inset shows the difference between the non-linear curve fits to the ABC (magenta, circles) and BAC (cyan, squares) data.

THIS PAGE IS INTENTIONALLY LEFT BLANK
